# Predicting behavior through dynamic modes in resting-state fMRI data

**DOI:** 10.1101/2021.05.22.445226

**Authors:** Shigeyuki Ikeda, Koki Kawano, Soichi Watanabe, Okito Yamashita, Yoshinobu Kawahara

**Affiliations:** RIKEN Center for Advanced Intelligence Project, Tokyo 103-0027 Japan; ATR Neural Information Analysis Laboratories, Kyoto 619-0288 Japan; Institute of Mathematics for Industry, Kyushu University, Fukuoka 819-0395 Japan

**Keywords:** Dynamic functional connectivity, Dynamic mode decomposition, Resting-state fMRI, Prediction, Behavior

## Abstract

Dynamic properties of resting-state functional connectivity (FC) provide rich information on brain-behavior relationships. Dynamic mode decomposition (DMD) has been used as a method to characterize FC dynamics. However, it remains unclear whether dynamic modes (DMs), spatial-temporal coherent patterns computed by DMD, provide information about individual behavioral differences. This study established a methodological approach to predict individual differences in behavior using DMs. Furthermore, we investigated the contribution of DMs within each of seven specific frequency bands (0-0.1,…,0.6–0.7 Hz) for prediction. To validate our approach, we confirmed whether each of 59 behavioral measures could be predicted by performing multivariate pattern analysis on a gram matrix, which was created using subject-specific DMs computed from resting-state functional magnetic resonance imaging (rs-fMRI) data of individuals. The prediction was successful, and DMD outperformed temporal independent component analysis, a conventional data decomposition method for extracting spatial activity patterns. Most of the behavioral measures that showed significant prediction accuracies in a permutation test were cognitive-behavioral measures. Our results suggested that DMs within frequency bands <0.2 Hz primarily contributed to prediction. In addition, we found that DMs <0.2 Hz had spatial structures similar to several common resting-state networks. We demonstrated the effectiveness of DMs, indicating that DMD is a key approach for extracting spatiotemporal features from rs-fMRI data.

## 1. Introduction

Resting-state brain activity is formed by spatially and temporally organized cortical networks (Greicius et al., 2003; Ven et al., 2004; Damoiseaux et al., 2006). Functional connectivity (FC) has been used extensively as a quantitative measure for characterizing the nature of interactions between cortical networks. It represents a temporal correlation between brain activities (Friston et al., 1993; Biswal et al., 1995). Previous studies have reported associations between FC and individual differences in behavior, e.g., intelligence (Song et al., 2008), emotional intelligence (Takeuchi et al., 2013), and personality (Adelstein et al., 2011), using large behavioral datasets (Smith et al., 2015). In addition to behavior, FC has been known to be associated with several psychiatric diseases, e.g., Alzheimer’s disease (Wang et al., 2007), major depression (Greicius et al., 2007), schizophrenia (Lynall et al., 2010; Skudlarski et al., 2010), and autism (Kennedy and Courchesne, 2008). FC provides useful information, but it is a static measure, invariant to temporal re-ordering of data points (Liégeois et al., 2017). Therefore, a new approach for characterizing dynamic FC has gained significant interest.

A sliding window approach has frequently been used to characterize dynamic FC (or time-varying FC) (Chang and Glover, 2010; Sakoğlu et al., 2010; Kiviniemi et al., 2011; Hutchison et al., 2013; Leonardi et al., 2013; Allen et al., 2014). In this approach, a temporal window with a constant length is prepared, and dynamic FC can be assessed by calculating FC over a time series of windowed segments. In addition, consistent spatial patterns can be extracted from dynamic FC using exploratory methods, such as clustering or principal component analysis, and these spatial patterns have been associated with behavior and neurodegenerative and psychiatric diseases (Wang et al., 2016; Leonardi et al., 2013; Damaraju et al., 2014). However, the choice of the window length remains a matter of debate (Preti et al., 2017). As an alternative to using a sliding window approach, a hidden Markov model (HMM) has also been used (Eavani et al., 2013; Baker et al., 2014), and consistent spatial patterns obtained by HMM have been reported to be associated with behavior (Vidaurre et al., 2017). However, there are no well-established methods for determining the number of hidden states for an HMM. Furthermore, because individual states are described using spatial patterns only, the HMM does not provide information about the temporal behavior of each spatial pattern. HMM with a multivariate autoregressive model (HMM-MAR) has been used to describe both spatial patterns that appear consistently in time and their temporal behavior, i.e., spectral properties (Vidaurre et al., 2016). HMM-MAR has two drawbacks: the number of MAR parameters escalates rapidly with an increase in hidden states; it is difficult to interpret the temporal behavior of individual hidden states because their spectral properties consist of many frequency components.

Dynamic mode decomposition (DMD), an algorithm originally developed in the field of fluid dynamics (Rowley et al., 2009; Schmid, 2010), has gained much attention as a method to characterize temporal behavior of brain activity. Because DMD can compute consistent spatial patterns that fluctuate with a single frequency, the temporal behavior of each spatial pattern is easily interpretable. Furthermore, DMD has almost no parameters that should be tuned. Pioneering studies have demonstrated the effectiveness of the method on electrocorticographic (ECoG) data and functional magnetic resonance imaging (fMRI) data (Brunton et al., 2016; Casorso et al., 2019; Shiraishi et al., 2020). Features computed by DMD can be expressed by both spatial and temporal characteristics inherent in the data. Specifically, DMD computes eigenvectors and corresponding eigenvalues of the approximate linear transformation expressing the time evolution of multidimensional time-series data. Eigenvectors (i.e., spatial characteristics) represent dynamic modes (DMs), which are coherent spatial structures, and the corresponding eigenvalues (i.e., temporal characteristics) represent the frequency and growth/decay rate in time. A previous study had applied DMD to resting-state brain activity measured by means of fMRI and investigated associations between temporal characteristics of DMD and human behavioral data (Casorso et al., 2019). Nevertheless, associations between the spatial characteristics of DMD and human behavioral data remain unclear because a quantitative method to assess associations between individual spatial characteristics and behavioral data has yet to be established.

The present study had three purposes: first, establishing a methodological approach in which individual differences of behavioral measures can be predicted using DMs computed by applying DMD to the resting-state fMRI data (rs-fMRI) of individuals; second, investigating which frequency band’s DMs contribute to prediction; and third, investigating whether computed DMs have spatial structures similar to common resting-state networks (RSNs).

As input features for predicting behavioral measures, we sought to compute a gram matrix representing the similarity between individuals using subject-specific DMs. However, the similarity cannot be computed by directly comparing DMs due to the lack of natural correspondence between subject-level DMs. Therefore, we employed the Grassmann manifold, on which a subspace spanned by a set of DMs can be represented as a point. The similarity between two sets of DMs is defined as the Frobenius norm between the two points on the Grassmann manifold (Hamm and Lee, 2008). In order to investigate the contribution of specific frequency bands to prediction, we computed a gram matrix using DMs within each of seven specific frequency bands (0–0.1,…, 0.6–0.7 Hz). After computing gram matrices, Gaussian process regression (GPR) was performed on the gram matrices to predict each of the 59 behavioral measures. These behavioral measures can cover seven behavioral categories (e.g., cognition). To validate our prediction method, we used behavioral and rs-fMRI data of the Human Connectome Project (HCP) dataset (Van Essen et al., 2013). Taken together, the present study demonstrated that behavioral measures could be predicted by our prediction method with spatial characteristics of DMD. Furthermore, we confirmed that DMD outperforms temporal independent component analysis (ICA), a conventional method to extract spatial activity patterns. Our results emphasize the effectiveness of DMD and suggest that DMD will become a promising approach to extract spatiotemporal features from rs-fMRI data.

## 2. Material and methods

### 2.1. Dataset

We used rs-fMRI data of the HCP 1200-subjects release (https://www.humanconnectome.org/study/hcp-young-adult/). After consulting with the Safety Management Division at RIKEN, we confirmed that institutional approval was not required for the use of the HCP dataset. Each functional image was acquired with a temporal resolution of 0.72 s and a 2-mm isotropic spatial resolution using a customized 3-T Siemens Skyra scanner (Smith et al., 2013). Individual subjects underwent four rs-fMRI runs of 14.4 min each (1200 frames per run), with eyes open, with a relaxed fixation on a projected bright cross-hair on a dark background. The rs-fMRI data were preprocessed with the HCP spatial and temporal preprocessing pipelines. Briefly, spatial preprocessing included spatial distortion correction, motion correction, and transformation to the Montreal Neurological Institute template space (Glasser et al., 2013). Temporal preprocessing included high-pass temporal filtering, ICA-FIX cleanup, and motion-related time-course removal (Smith et al., 2013). Details on the ICA-FIX cleanup have previously been described (Salimi-Khorshidi et al., 2014; Griffanti et al., 2014). Our analysis began with the preprocessed rs-fMRI data (e.g., *rfMRI_REST1_LR_hp2000_clean.nii.gz*).

We selected 59 behavioral measures from the HCP dataset, which cover seven behavioral categories, i.e., cognition, alertness, motor, sensory, in-scanner task performance, personality, and emotion (Supplementary Table 1). These behavioral measures were comprised of 58 measures used in a previous study (Liégeois et al., 2019) and a measure (i.e., *CogFluid-Comp_Unadj).* In addition to the 59 measures, we used five subject demographics, i.e., age *(Age_in_Yrs),* gender (*Gender*), race (*Race*), education *(SSAGA_Educ),* and percentage of rs-fMRI scan completion *(3T_RS-fMRI_PctCompl).*

The HCP 1200-subjects release includes data on 1206 healthy young adult subjects. For our analysis, we selected subjects who met the following conditions: (i) Subjects with no missing data for the 59 behavioral measures and five subject demographics; (ii) Subjects who completed all the rs-fMRIruns (i.e., *3T_RS-fMRI_PctCompl);* (iii) Subjects who scored ≥27 on the Mini-Mental State Examination (i.e., *MMSE_Score)* (O’Bryant et al., 2008); (iv) Subjects who clearly reported their race (i.e., we excluded subjects with “Unknown or Not Reported”); and (v) Subjects with head motion ≤0.15 in all the rs-fMRI runs (i.e., *Movement_RelativeRMS_mean.txt).* Consequently, 829 subjects (388 men, 441 women, mean age: 28.6 ± 3.7 years) were selected based on the above conditions.

### 2.2. Data preprocessing

To remove confounding factors, age, gender, race, education, and head motion, averaged across all the runs, were regressed out from each behavioral measure using ordinary least-squares regression (Liégeois et al., 2019).

For each run for each subject, a mean functional image was computed by averaging all functional images, which was subtracted from individual functional images. Each of the subsequent functional images was parcellated into 268 nodes based on a predefined functional atlas (Shen et al., 2013; Finn et al., 2015). A time series for each node was calculated by averaging the voxel-wise time series within each node, resulting in a 268 × 1200 data matrix (i.e., R1,…, R4 in Figure 1).

**Figure 1:**
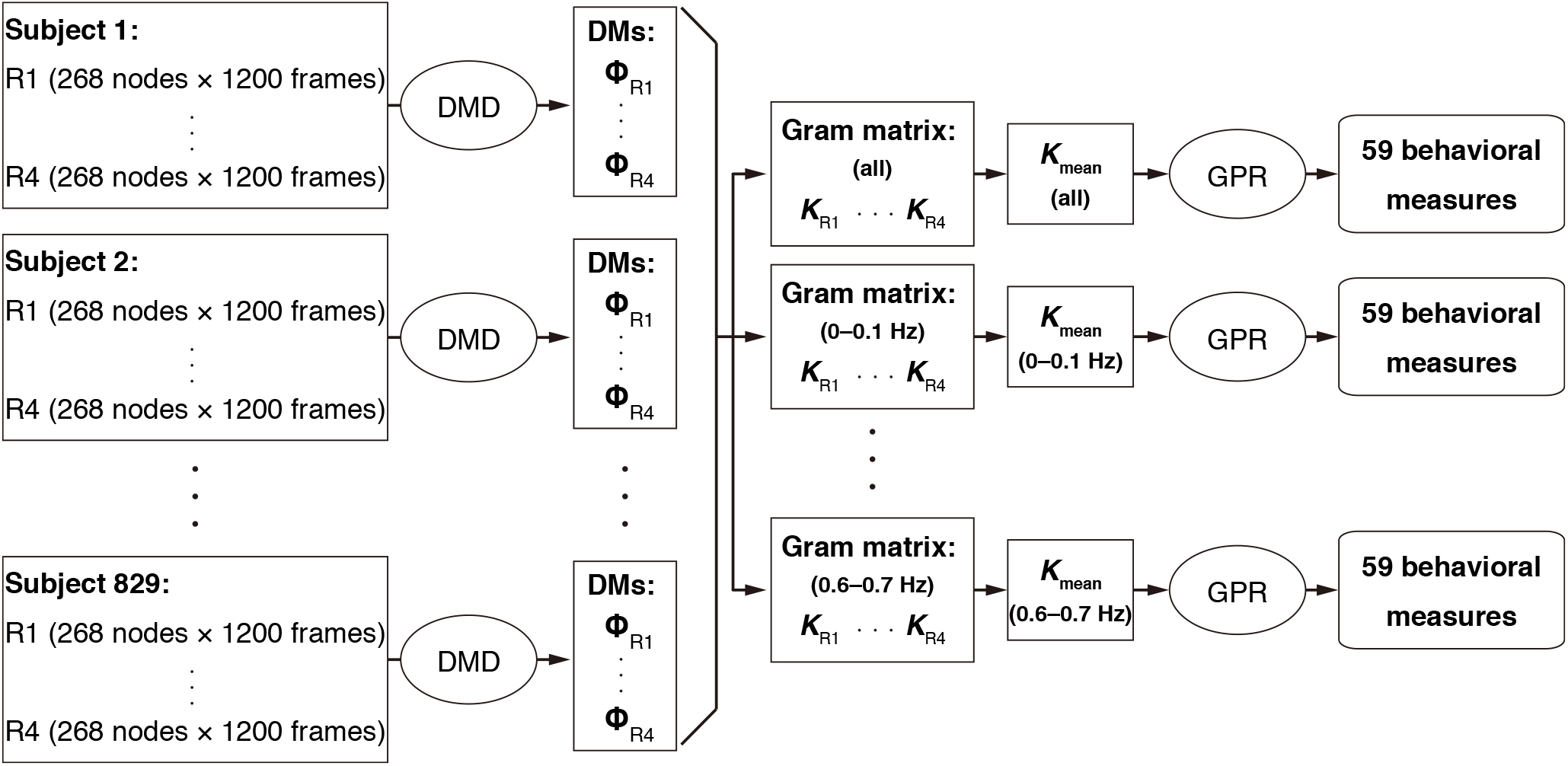
Graphical overview of the analysis procedure. Data for individual subjects were acquired in four resting-state fMRI runs. Using a functional atlas previously defined (Shen et al., 2013; Finn et al., 2015), each functional image was parcellated into 268 nodes and a time course for each node was calculated. The data for each run (i.e., R1, …, R4) were decomposed into dynamic modes (i.e., DMs: **Φ**_R1_, …, **Φ**_R4_) using dynamic mode decomposition (DMD). A gram matrix (i.e., ***K***_R1_, …, ***K***_R4_) was created using DMs within each specific frequency band (i.e., 0–0.1,…, 0.6–0.7 Hz). Note that “all” includes all frequency bands. A ***K***_meaa_ was calculated by averaging all the gram matrices in each specific frequency band. Each of the 59 behavioral measures was predicted using a mean gram matrix and Gaussian process regression (GPR).

### 2.3. DMD

Here, we outlined the exact DMD used in the present study. For more details, please refer to the previous study (Tu et al., 2014).

The fMRI time series from *n* nodes sampled every *k*Δ*t* can be described as follows: (***x***_1_, ***x***_2_, …, ***x**_m_* ∈ ℝ^*n*^), where Δ*t* represents the temporal resolution of rs-fMRI (= 0.72 s) and *m* represents the number of frames (= 1200). For example, ***x***_*k*_ is a spatial activity pattern sampled at *k*Δ*t*.

Now, two (*n* × *m* – 1) data matrices are created from the rs-fMRI data for each run for each subject (e.g., R1 in Figure 1):

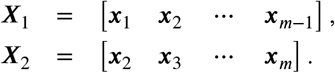

Suppose that we have an unknown linear transformation ***A***, such that

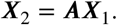

In general, ***x**_k_* can be enormously high-dimensional. The exact DMD of the pair ***X***_1_ and ***X***_2_ is given by computing the low-rank eigen-decomposition of ***A***, resulting in an exact eigenvector 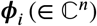 and eigenvalue 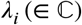. Each eigenvector *φ_i_* represents a DM (i.e., a coherent spatial structure) and the corresponding eigenvalue *λ_i_* represents its temporal characterization (i.e., frequency and growth/decay). Consequently, an approximation of the measured rs-fMRI data can be described as an underlying dynamic model:

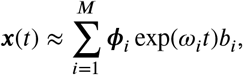

where *M* is the number of eigenvectors (or eigenvalues), *ω_i_* = ln (*Δ_i_*)/ Δ*t*, *t* is time, and ***b*** = **Φ**^†^***x***_1_.

The number of DMs given by DMD is the smaller of *n* and *m* – 1. If DMD is performed on the pair ***X***_1_ and ***X***_2_, we obtain *n* DMs because of *n* < *m* – 1. Generally, there are too few to capture the dynamics over the entire sampling time. To avoid this problem, we increased the rank of ***X***_1_ and ***X***_2_ by constructing a new augmented data matrix (Brunton et al., 2016)

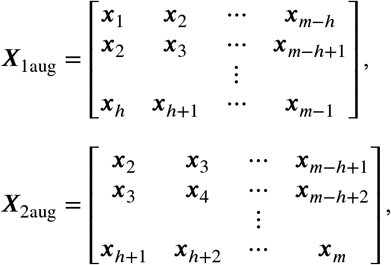

where *h* is the number of times that ***x***_*k*_ is stacked vertically. We chose *h* = 5, which is the smallest value, such that *hn* exceeds *m – h*. Consequently, DMD was performed on the pair ***X***_1aug_ and ***X***_2aug_ in the present study. In this way, we obtained a set of DMs and corresponding temporal characteristics for each run for each subject (Figure 1):

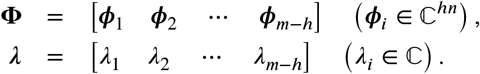

### 2.4. Construction of a gram matrix

As input features for prediction, we constructed a gram matrix representing the similarity of DMs between subjects. However, we could not simply compute the similarity of DMs using a common similarity index, such as the cosine similarity, because there is no natural correspondence between subject-level DMs. To overcome this problem, we developed a positive definite kernel *k* (**Φ**_*i*_, **Φ**_*j*_ referring to the projection kernel (Hamm and Lee, 2008), which corresponds to the metric on the Grassmann manifold:

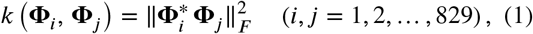

where **Φ** represents a set of DMs of individual subjects, || · ||_*F*_ represents the Frobenius norm, and the asterisk denotes Hermitian transpose. A subspace spanned by **Φ**_*i*_ can be represented as a point on the Grassmann manifold. As such, similarity between two subspaces (i.e., **Φ**_*i*_ and **Φ**_*j*_) can be computed by Eq. 1. Note that although the original projection kernel requires orthonormalization of **Φ**, each **Φ** is not orthonormalized in our kernel. Our kernel was designed to reflect the magnitudes of DMs to the projection kernel by weighting axes in the subspaces. Using the kernel, a gram matrix was computed from each run (e.g., ***K***_R1_ in Figure 1). Subsequently, a mean gram matrix (e.g., ***K***_mean_) was computed by averaging four gram matrices.

To investigate which frequency band’s DMs are important for prediction, we constructed a gram matrix using DMs within a specific frequency band. The whole frequency spectrum in the rs-fMRI data reached 0.69 Hz (Nyquist frequency). The frequency spectrum was divided by intervals of 0.1 Hz so that seven specific frequency bands (i.e., 0–0.1,…,0.6–0.7 Hz) were obtained.

### 2.5. Prediction analysis

We used GPR to predict individual differences of each of the 59 behavioral measures from the mean gram matrix (Figure 1). The subjects in the HCP dataset included siblings. To prevent the family structure from biasing prediction results, we employed a leave-one-family-out cross-validation procedure (Dubois et al., 2018a,b). Prediction accuracy was estimated by calculating the Pearson correlation coefficient and root mean squared error (RMSE) between the predicted and true behavioral scores. To compute *p* values of both a correlation coefficient and an RMSE value, a permutation test was performed (10,000 times). In particular, we randomly permuted all behavioral scores but preserved the structure of the mean gram matrix and performed the leave-one-family-out cross-validation procedure. A *p* value corresponding to the true correlation was defined as the probability of observing a correlation equal to or greater than the true correlation. In addition, a *p* value corresponding to the true RMSE value was defined as the probability of observing an RMSE value equal to or less than the true RMSE value. Consequently, we obtained 59 *p* values for each correlation and RMSE. To test the statistical significance of prediction accuracy, we applied a false discovery rate correction (*q* ≤ 0.05) to each correlation and RMSE (Benjamini and Hochberg, 1995). The prediction accuracy was regarded as significant only if both the correlation and RMSE were significant.

### 2.6. Temporal ICA

For comparison, we performed prediction using temporal ICA. We chose temporal ICA rather than spatial ICA because there were fewer spatial dimensions of the data (268) than temporal dimensions (1200). Before temporal ICA, global scaling was applied to data for each run (e.g., R1 in Figure 1). Specifically, the individual node time-series was demeaned and divided by the global standard deviation, the mean value of all the standard deviations of the nodes. The subsequent data were decomposed into independent components using FastICA2.5 (Hyvärinen, 1999) (https://research.ics.aalto.fi/ica/fastica/code/dlcode.shtml):

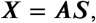

where ***X*** = [***x***_1_ ***x***_2_ … ***x**_m_*], ***A*** is the mixing matrix, in which each column represents a spatial map, and ***S*** is the separating matrix, in which each row represents a temporal independent component. For each run, a gram matrix was constructed from the spatial maps ***A*** in the same manner as for DMD. Subsequently, a mean gram matrix was computed by averaging four gram matrices.

### 2.7. Clustering of DMs

We used the modified K-means to summarize entire DMs (Pascual-Marqui et al., 1995). An individual DM ***ϕ***_*i*_ was a *hn*×1 vector because the spatial dimension of the augmented data matrix was *hn*; ***ϕ***_*i*_ is *h* stacks of essentially identical repeats (Brunton et al., 2016). Therefore, we used the first *n* elements of individual DMs in the clustering analysis.

Because DMs are complex-valued, we could not apply the modified K-means to them. To avoid this problem, we created a vector (∈ ℝ^2*n*^) containing the real and imaginary parts of each DM and aggregated all vectors (i.e., all the runs for all the subjects) into a matrix. We created an aggregate matrix for each specific frequency band and applied the modified K-means to individual aggregate matrices. Note that cluster centroids were normalized such that they are of unit vector (see Pascual-Marqui et al. (1995), for details).

The optimum number of cluster centroids was determined using the cross-validation criterion (Pascual-Marqui et al., 1995). Specifically, we used the modified K-means while varying the number of clusters (2–15) and chose a value that minimized the cross-validation criterion. Finally, using the real and imaginary parts of centroids, magnitude and phase maps were computed separately.

We quantitatively assessed how strongly the cluster centroids were related to common RSNs. Specifically, the cosine similarity was calculated between each magnitude map and each of the 10 previously defined RSNs (Smith et al., 2009) and between each phase map and each of the RSNs.

## 3. Results

### 3.1. DMD vs. temporal ICA

We investigated whether individual differences of behavioral measures could be predicted from DMs and whether DMD yielded better predictive power than temporal ICA. DMD showed significant prediction accuracies in seven behavioral measures, i.e., *CardSort_Unadj, Flanker_Unadj, Emotion_Task_Face_Acc, CogFluidComp_Unadj*, *WM_Task_Acc, ReadEng_Unadj,* and *ER40ANG* (Figure 2 and Supplementary Table 2). Scatter plots between predicted and true behavioral scores of individual behavioral measures are shown in Supplementary Figure 1. These results indicated that individual differences in the behavioral measures could be predicted from DMs. Temporal ICA showed significant prediction accuracies in three behavioral measures, i.e., *WM_Task_Acc, ReadEng_Unadj*, and *NEOFAC_E*; two of these significant measures were also significant in DMD. Although we did not statistically compare prediction accuracies between DMD and temporal ICA due to not being able to estimate the variability of each prediction accuracy, DMD was found to outperform temporal ICA. On the other hand, we did not observe significant prediction accuracies in many behavioral measures.

**Figure 2:**
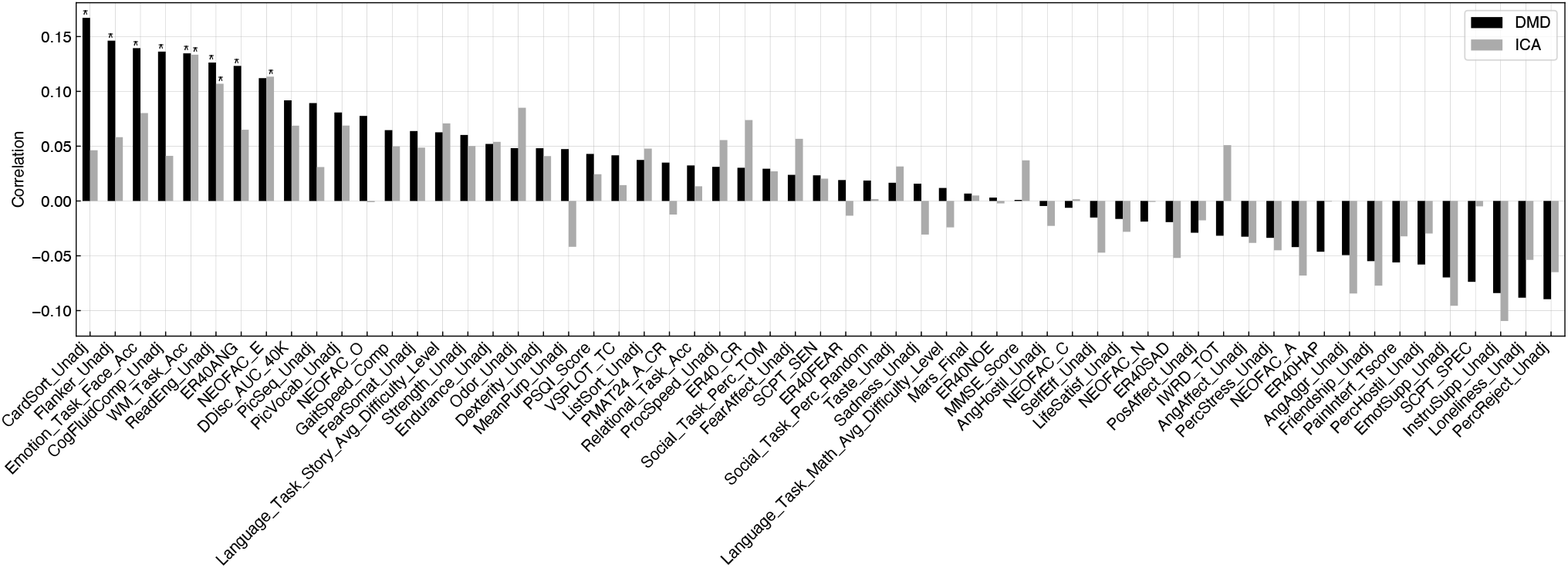
Comparison of prediction results between dynamic mode decomposition (DMD) and temporal independent component analysis (ICA). Prediction accuracy (Pearson correlation coefficient) for each of the 59 behavioral measures is presented. Asterisks indicate prediction accuracies that were significant in both the correlation and root mean squared error (*q* < 0.05).

### 3.2. Prediction results in different frequency bands

We investigated which frequency band’s DMs were important in prediction and into which categories significant behavioral measures fell. First, we found that the use of all the frequency bands provided better predictive power than the use of individual-specific frequency bands (Figure 3). Most of the significant prediction accuracies were observed in the cognition category, i.e., *CardSort_Unadj, Flanker_Unadj, CogFluidComp_Unadj,* and *ReadEng_Unadj.* Frequency bands <0.2 Hz showed some significant prediction results, which were also significant in “all”. On the other hand, no significant prediction accuracies, except for two measures (i.e., *Endurance_Unadj* and *Taste_Unadj),* were observed in high-frequency bands (≥ 0.2 Hz).

**Figure 3:**
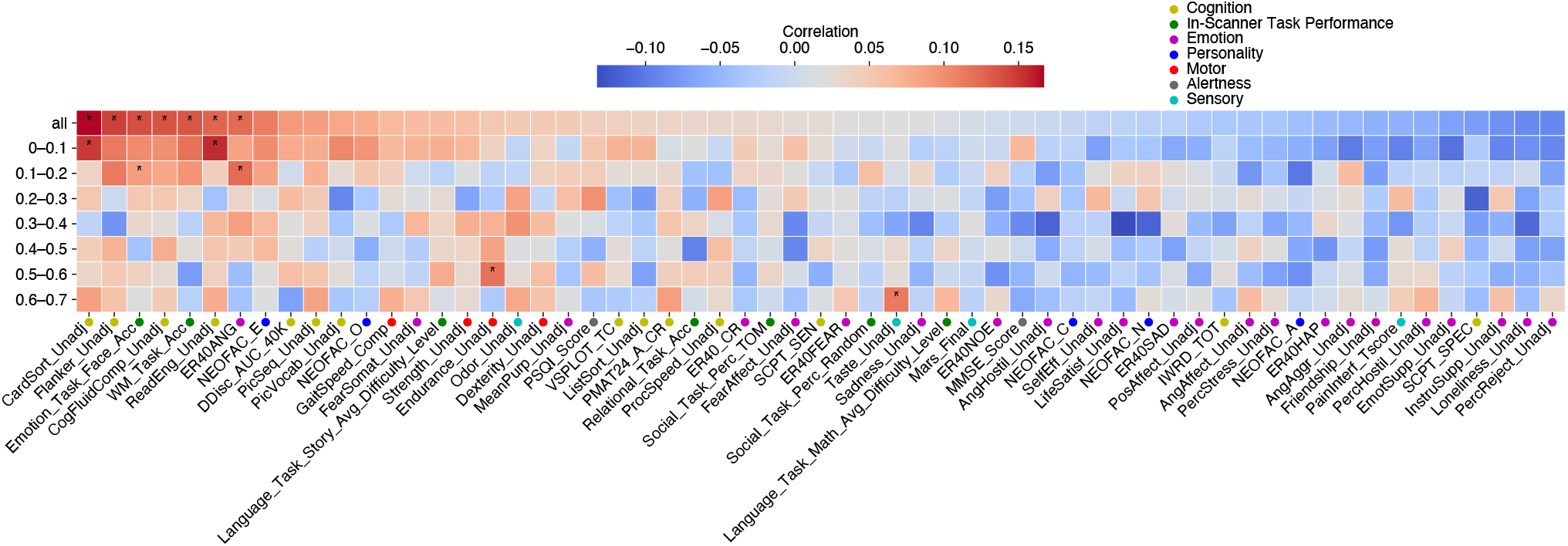
Comparison of prediction results among different frequency bands. Prediction accuracies (Pearson correlation coefficient) for each frequency band are presented. Note that “all” indicates prediction results obtained by using all the frequency bands, so that the results are the same as those of the dynamic mode decomposition shown in Figure 2. Asterisks indicate prediction accuracies that were significant in both the correlation and root mean squared error (*q* ≤ 0.05). All the behavioral measures were classified into seven categories, each of which was colored.

### 3.3. Cluster centroids of DMs

We computed cluster centroids of DMs obtained from all the runs for all the subjects and investigated the relevance of the centroids to the common RSNs. We chose the optimum number of clusters for each frequency band based on the cross-validation criterion (Supplementary Figure 2). The optimum number of clusters in the 0–0.1 Hz band was ten, and those in the other bands were two or three.

Magnitude and phase maps of 10 centroids in the 0–0.1 Hz band are displayed in Figure 4a. Cosine similarities of these maps with the 10 common RSNs (Smith et al., 2009) are displayed in Figure 4b. In magnitude, the medial visual area (Med-Vis) was strongly associated with centroids 3, 5, 7, and 8. The lateral visual area (Lat-Vis) was moderately related to centroid 3. The default mode network (DMN) was associated with centroids 9 and 10. The executive control (EC) showed associations with centroids 1, 2, 6, and 9. The left frontoparietal area (L-FP) was moderately related to centroid 6. On the other hand, in-phase, the DMN showed associations with centroids 1 and 10. The sensorimotor area (SM) was related to centroid 6. The EC was associated with centroid 3.

**Figure 4:**
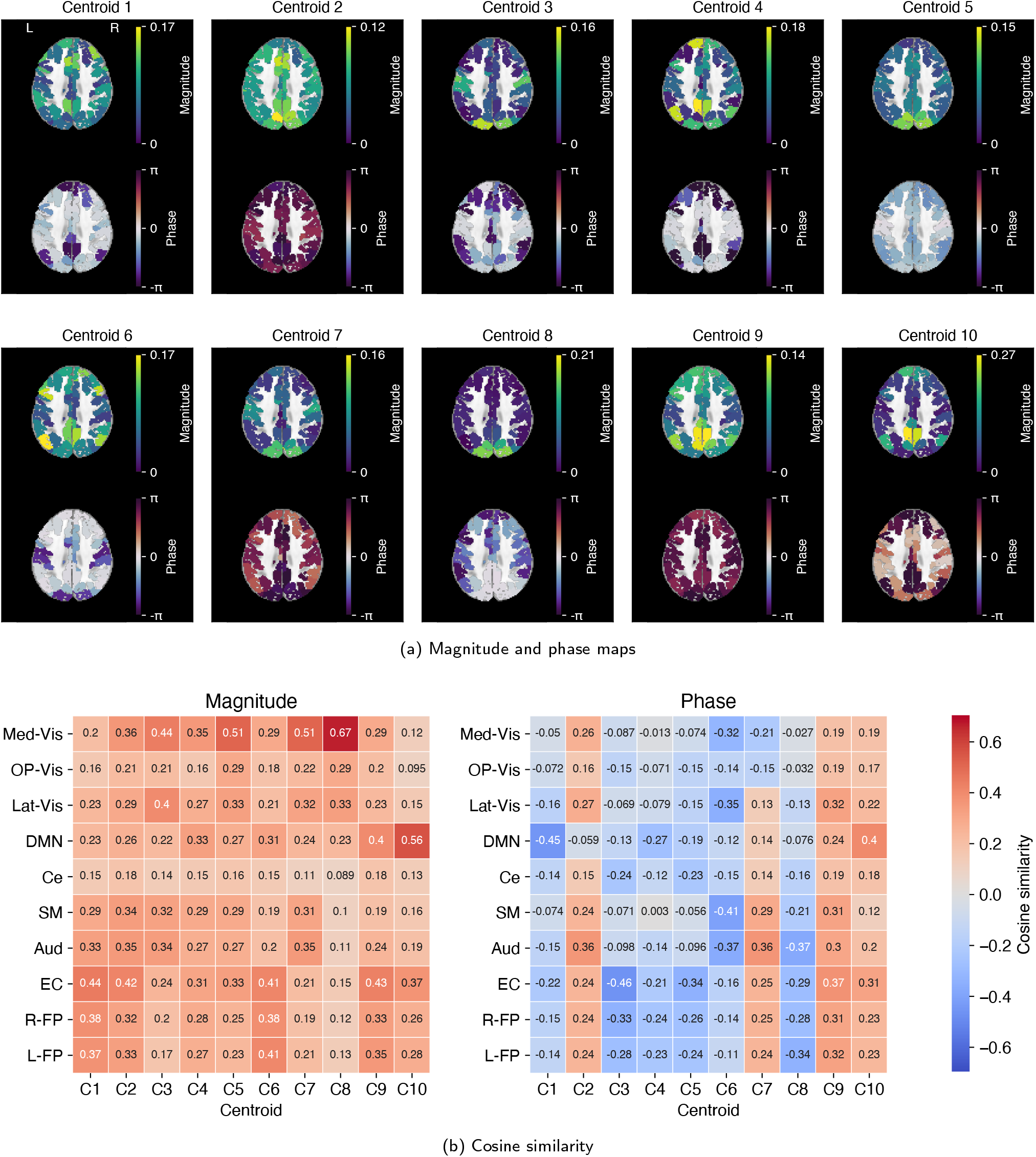
Clustering results (0–0.1 Hz). (a) Magnitude and phase maps of 10 centroids. An axial image (z = 38) for each map is shown superimposed on the skull-stripped version of the Montreal Neurological Institute (MNI152) standard space template image. (b) Cosine similarities between magnitude maps and the 10 resting-state networks (RSNs) and between phase maps and the RSNs. The 10 RSNs are as follows: the medial, occipital pole, and lateral visual areas (Med-Vis, OP-Vis, and Lat-Vis), the default mode network (DMN), the cerebellum (Ce), the sensorimotor area (SM), the auditory area (Aud), the executive control (EC), and the right and left frontoparietal areas (R-FP and L-FP).

Clustering results in the other frequency bands are displayed in Supplementary Figures 3–8. In the magnitude of the 0.1–0.2 Hz band, the Med-Vis was strongly associated with centroids 2 and 3, and the EC was related to centroid 1. In the magnitude of the frequency bands (≥ 0.2 Hz), only the EC was related to several centroids from multi-frequency bands. In-phase, almost no centroids showed associations with the RSNs.

## 4. Discussion

The main goal of the present study was to establish a methodological approach to predict individual differences of human behaviors from the spatial characteristics of DMD (i.e., DMs). Fifty-nine behavioral measures covering seven behavioral categories were chosen from the HCP dataset, and individual differences of each behavioral measure were predicted using DMs computed from the rs-fMRI data. We observed significant prediction accuracies in seven of these behavioral measures, indicating that the prediction was successful. A previous study had already clarified the associations between temporal characteristics of DMD (i.e., eigenvalues) and behavioral measures using canonical correlation analysis (Casorso et al., 2019). Therefore, our study provides an understanding of the link between the properties of resting-state DMs and human behaviors. Another noteworthy difference between the present study and Casorso et al. (2019) is that while Casorso et al. (2019) employed a correlational approach, we employed a prediction approach. We evaluated the generalizability of DMs-behavior relationships. For comparison, we used temporal ICA and found that DMD outperformed temporal ICA. These results suggest that DMs, spatial-temporal coherent patterns, yield better predictive power than spatial activity patterns computed by temporal ICA. Furthermore, the prediction analysis with specific frequency bands suggests that DMs <0.2 Hz primarily contribute to prediction. In addition, we found that DMs had spatial structures similar to the RSNs, suggesting that the well-known RSNs (e.g., the DMN) have coherent dynamics in the resting-state brain activity. Our study highlighted the importance of the data decomposition approach for extracting consistent brain activity patterns in both space and time.

### 4.1. Prediction results in DMD

Across all 59 behavioral measures (Figure 2), DMD showed an average prediction accuracy of *correlation* = 0.0233, and ICA showed an average prediction accuracy of *correlation* = 0.0115. HCP MegaTrawl (https://db.humanconnectome.org/megatrawl/) predicted HCP behavioral measures using functional connectivity (i.e., partial correlation) computed from rs-fMRI data. In the HCP MegaTrawl, prediction accuracies for 48 of 59 behavioral measures were reported. Across the 48 behavioral measures, the HCP MegaTrawl showed an average prediction accuracy of correlation = −0.0254 (deconfounded space). Since the HCP MegaTrawl employed a different methodological approach from our own (e.g., removing confounding factors, regression model, and crossvalidation procedure), these average prediction accuracies could not be simply compared. Nevertheless, DMD seems to be comparable to typical functional connectivity.

### 4.2. Spatial-temporal coherent patterns

We found that DMD showed better predictive power than temporal ICA. Temporal ICA decomposes rs-fMRI data into spatial components that are temporally independent of each other, while DMD decomposes it into spatial components that have coherent dynamics with a specific frequency. Our results indicated the importance of spatial-temporal coherent patterns in the resting-state brain activity and supported our previous study (Shiraishi et al., 2020). That study investigated whether three types of hand movement could be classified using DMs computed from ECoG signals and compared classification accuracies between DMD and fast Fourier transform (FFT). DMD outperformed FFT. Taken together with the current study results, this suggests that spatial activity patterns with coherent dynamics encode a variety of information, ranging from motor function to higher-order cognitive function.

### 4.3. Grassmann manifold

We used the Grassmann manifold to facilitate direct comparison of subject-specific DMs. The use of the Grassmann manifold led to the successful prediction of behavioral measures. The Grassmann manifold has been used in a pioneering study in the field of image identification (Hamm and Lee, 2008). In the neuroscience field, a study used the Grassmann manifold to compare individual-specific independent components computed from rs-fMRI data and succeeded in distinguishing patients with schizophrenia from healthy controls (Fan et al., 2011). Our recent study adopted the Grassmann manifold to compute the similarity between sets of DMs and indicated its effectiveness (Shiraishi et al., 2020). Therefore, our prediction results supported these previous findings. On the other hand, a previous study avoided the problem of the lack of natural correspondence between individual DMs by computing group DMs (Casorso et al., 2019). In particular, the previous study imposed subject-specific DMs to be equal to the group DMs and obtained subject-specific temporal characteristics corresponding to the group DMs. Although the approach is valid, this requires the calculation of group DMs. The Grassmann manifold allows direct comparison of individual DMs with no additional computational cost.

### 4.4. Categories of significant behavioral measures

For prediction results obtained using all frequency bands (i.e., “all” in Figure 3), most of the significant prediction accuracies were observed in the cognition category. On the other hand, only one significant prediction accuracy (i.e., *ER40ANG*) was observed in the emotion category. The results are partially consistent with those of a previous study that performed a canonical correlation analysis between temporal characteristics of DMD computed from rs-fMRI data and 158 HCP behavioral and demographic subject measures (Casorso et al., 2019). Specifically, the previous study found two significant canonical modes linking temporal characteristics and subject measures, and the modes correlated with 27 subject measures. Note that some subject measures correlated with both the modes. The cognition category was the most common category into which the 27 measures fell, followed by the emotion category. Although it is difficult to make a direct comparison between our own and the previous results because of the use of different methodological approaches, the comparison has an intriguing implication, that is, both spatial and temporal characteristics of rs-fMRI data encode information about individual differences in behavioral measures in the cognition category, while, unlike temporal characteristics, spatial characteristics moderately encode information about individual differences in emotion category measures.

### 4.5. Contribution of specific frequency bands to prediction

Using all the frequency bands (i.e., “all” in Figure 3) was found to provide better predictive power than using a specific frequency band. This suggests that DMs from multifrequency bands contain complementary information about individual differences among multiple frequency bands. In fact, combining biomarkers from multi-frequency bands has been found to improve the classification accuracy of schizophrenia (Huang et al., 2019).

We found some significant prediction accuracies in frequency bands <0.2 Hz, which were also significant in “all.” These results suggest that prediction results of “all” primarily arise from DMs <0.2 Hz. In comparison with the 0–0.1 Hz band (the typical band for rs-fMRI), the 0.1–0.2 Hz band has attracted less attention in previous rs-fMRI research. However, several previous studies on rs-fMRI have reported intriguing findings: stronger spectral power (approximate 0.12– 0.2 Hz) in patients with chronic pain than in control subjects (Malinen et al., 2010; Baliki et al., 2011), a relationship between the slow-3 band (0.073–0.198 Hz) and personality (Ikeda et al., 2017), differences in amplitude of the slow-3 band between healthy subjects and those with psychosis (Gohel et al., 2018), and the moderately high classification accuracy of schizophrenia based on the slow-3 band (Huang et al., 2019). These previous findings have indicated the importance of the 0.1–0.2 Hz band in rs-fMRI research, which was supported by our present results. On the other hand, we found only two significant prediction accuracies in higher frequency bands (≥ 0.2 Hz), which were not significant in “all.” Although the higher frequency bands seem to provide less predictive power than the lower frequency bands (< 0.2 Hz), they may encode information about individual differences that differ from those encoded by the lower frequency bands. Together, this study emphasizes the need for further investigation of not only the typical band but also frequency bands >0.1 Hz in rs-fMRI.

### 4.6. Relations between DMs and RSNs

For frequency bands <0.2 Hz, most magnitude maps of centroids were found to relate to the five common RSNs (i.e., 0–0.1 Hz: Med-Vis, Lat-Vis, DMN, EC, and L-FP; 0.1–0.2 Hz: Med-Vis and EC). A pioneering study that applied DMD to rs-fMRI data computed representative DMs and found that the representative DMs had spatial structures similar to the DMN and visual area (Casorso et al., 2019). For frequency bands of ≥0.2 Hz, intriguingly, only the EC had associations with centroids from multi-frequency bands. These results suggest that many representative DMs in the lower frequency bands (< 0.2 Hz) but not those in the higher frequency bands (≥ 0.2 Hz) had spatial structures similar to common RSNs. This supports our implication that DMs <0.2 Hz contribute to prediction. The range of 0–0.1 Hz is a well-known frequency band in rs-fMRI research. Actually, a previous study reported that the major frequency contributions to resting-state FC resulted from fluctuations <0.1 Hz (Cordes et al., 2001). Furthermore, RSNs have also been observed in the 0.1–0.2 Hz band (Wu et al., 2008; Salvador et al., 2008; Baria et al., 2011; Niazy et al., 2011; Gohel and Biswal, 2015; Wang et al., 2018,2020). These previous findings are consistent with our results. However, other previous studies found RSNs in frequency bands of >0.2 Hz (Lee et al., 2013; Boubela et al., 2013; Gohel and Biswal, 2015; Wang et al., 2020). The inconsistency between the previous results and our results may imply that it is difficult to extract DMs that have a spatial structure similar to those of RSNs in the high-frequency range. A previous rs-fMRI study investigated time-frequency dynamics of coherence between different regions within the DMN and found that the coherence tends to decrease with an increase in the fMRI signal frequency (Chang and Glover, 2010). The previous findings suggested that, with an increase in fMRI signal frequency, it is difficult to compute spatial activity patterns that have coherent dynamics (i.e., DMs) from rs-fMRI data. This may explain the inconsistent results.

For the 0–0.1 Hz band, phase maps of centroids showed associations with some RSNs (i.e., the DMN, SM, and EC). Because many previous rs-fMRI studies have discovered RSNs using Pearson correlation coefficient, it is reasonable to think that brain regions constituting RSNs fluctuate synchronously (i.e., phase coherence). Phase maps in the 0.1-0.2 Hz band did not show associations with RSNs, even though magnitude maps in the frequency band were related to some RSNs. Furthermore, for the higher frequency bands (≥ 0.2), almost no phase maps were related to RSNs. It seems that these results may be explained by previous findings (Chang and Glover, 2010) that showed that the coherence between regions of RSNs tends to decrease with an increase in fMRI signal frequency.

### 4.7. Limitations

The present study had some limitations. First, we used only spatial characteristics of DMD in order to predict individual differences of behavioral measures. As shown by a previous study, temporal characteristics of DMD also contain information about individual differences in various human behaviors (Casorso et al., 2019). A previous study proposed kernels containing information about both spatial and temporal characteristics (e.g., Koopman trace kernel) (Fujii et al., 2017). It is deemed desirable to compute a gram matrix using the proposed kernels. Second, although DMD can be applied to rs-fMRI data without using parcellation, we employed it due to considerations of the computational cost. Third, the present study used only rs-fMRI data of the HCP dataset. Previous studies demonstrated the difficulties of using multi-site rs-fMRI data (Turner et al., 2013; Nielsen et al., 2013). Further investigation of the effectiveness of DMD on rs-fMRI data from multi-sites is needed. Fourth, we used cosine similarity to evaluate the similarity between phase maps and RSNs. The phase is cyclic; *π* and *-π* are different values, but are the same in terms of phase. On the other hand, RSNs were all ICA spatial maps. This raises doubt about whether the similarity between phase maps and RSNs can be computed using cosine similarity. However, at present, there is no valid alternative for computing similarity between phase maps and RSNs.

## 5. Conclusions

The present study succeeded in establishing the methodological approach to predicting individual differences in behavioral measures from DMs that are inherent in rs-fMRI activity. It indicated that DMD outperformed temporal ICA. The prediction analysis based on specific frequency bands suggested that DMs <0.2 Hz largely contribute to the prediction. This was supported by the clustering analysis, in which we found many spatial structures similar to RSNs from magnitude maps of <0.2 Hz but not those ≥ 0.2 Hz. Furthermore, DMs may primarily encode information about individual differences in cognitive-behavioral measures. Taken together, our findings emphasize the superiority of DMD in the data decomposition approach for extracting spatiotemporal features from rs-fMRI data. Future studies should investigate whether DMs can be used as a biomarker for psychiatric diseases, which will facilitate a deeper understanding of the effectiveness of DMD.

## 6. Acknowledgments

This work was supported in part by JSPS KAKENHI Grant Numbers JP18H03287,JP19K20311, the Cooperative Research Project Program of Joint Usage/Research Center at the Institute of Development, Aging and Cancer, Tohoku University, AMED under Grant Number JP20dm0307009, and JST, CREST Grant Number JPMJCR1913, Japan. Data were provided by the Human Connectome Project, WU-Minn Consortium (Principal Investigators: David Van Essen and Kamil Ugurbil; 1U54MH091657) funded by the 16 NIH Institutes and Centers that support the NIH Blueprint for Neuroscience Research; and by the McDonnell Center for Systems Neuroscience at Washington University.

## CRediT authorship contribution statement

**Shigeyuki Ikeda:** Software, Formal analysis, Writing – Original Draft, Visualization, Funding acquisition. **Koki Kawano:** Software, Formal analysis, Writing – Review & Editing. **Soichi Watanabe:** Software, Formal analysis, Writing – Review & Editing. **Okito Yamashita:** Conceptualization, Writing – Review & Editing, Supervision, Funding acquisition. **Yoshinobu Kawahara:** Conceptualization, Methodology, Writing – Review & Editing, Supervision, Funding acquisition.

