## Supplementary materials for "Predicting behavior through dynamic modes in resting-state fMRI data"

Supplementary Table 1: List of the 59 behavioral measures selected from the Human Connectome Project dataset. The measures can cover seven behavioral categories, i.e., Cognition, Alertness, Motor, Sensory, In-Scanner Task Performance, Personality, and Emotion. Please refer to *"HCP_S1200_DataDictionary_April_20_2018.csv"*.

| | columnHeader | fullDisplayName | category |
|---|---|---|---|
| 1 | CogFluidComp_Unadj | NIH Toolbox Cognition Fluid Composite: Unadjusted Scale Score | Cognition |
| 2 | PicSeq_Unadj | NIH Toolbox Picture Sequence Memory Test: Unadjusted Scale Score | Cognition |
| 3 | CardSort_Unadj | NIH Toolbox Dimensional Change Card Sort Test: Unadjusted Scale Score | Cognition |
| 4 | Flanker_Unadj | NIH Toolbox Flanker Inhibitory Control and Attention Test: Unadjusted Scale Score | Cognition |
| 5 | PMAT24_A_CR | Penn Progressive Matrices: Number of Correct Responses (PMAT24_A_CR) | Cognition |
| 6 | ReadEng_Unadj | NIH Toolbox Oral Reading Recognition Test: Unadjusted Scale Score | Cognition |
| 7 | PicVocab_Unadj | NIH Toolbox Picture Vocabulary Test: Unadjusted Scale Score | Cognition |
| 8 | ProcSpeed_Unadj | NIH Toolbox Pattern Comparison Processing Speed Test: Unadjusted Scale Score | Cognition |
| 9 | DDisc_AUC_40K | Delay Discounting: Area Under the Curve for Discounting of $40,000 (DDisc_AUC_40K) | Cognition |
| 10 | VSPLOT_TC | Variable Short Penn Line Orientation: Total Number Correct (VSPLOT_TC) | Cognition |
| 11 | SCPT_SEN | Short Penn Continuous Performance Test: Sensitivity = SCPT_TP/(SCPT_TP + SCPT_FN) (SCPT_SEN) | Cognition |
| 12 | SCPT_SPEC | Short Penn Continuous Performance Test: Specificity = SCPT_TN/(SCPT_TN + SCPT_FP) (SCPT_SPEC) | Cognition |
| 13 | IWRD_TOT | Penn Word Memory Test: Total Number of Correct Responses (IWRD_TOT) | Cognition |
| 14 | ListSort_Unadj | NIH Toolbox List Sorting Working Memory Test: Unadjusted Scale Score | Cognition |
| 15 | MMSE_Score | Mini Mental Status Exam Total Score | Alertness |
| 16 | PSQI_Score | Sleep (Pittsburgh Sleep Questionnaire) Total Score | Alertness |
| 17 | Endurance_Unadj | NIH Toolbox 2-minute Walk Endurance Test : Unadjusted Scale Score | Motor |
| 18 | GaitSpeed_Comp | NIH Toolbox 4-Meter Walk Gait Speed Test: Computed Score | Motor |
| 19 | Dexterity_Unadj | NIH Toolbox 9-hole Pegboard Dexterity Test : Unadjusted Scale Score | Motor |
| 20 | Strength_Unadj | NIH Toolbox Grip Strength Test: Unadjusted Scale Score | Motor |
| 21 | Odor_Unadj | NIH Toolbox Odor Identification Age 3+ Unadjusted Scale Score | Sensory |
| 22 | PainInterf_Tscore | NIH Toolbox Pain Interference Survey Age 18+: T-score | Sensory |
| 23 | Taste_Unadj | NIH Toolbox Regional Taste Intensity Age 12+ Unadjusted Scale Score | Sensory |
| 24 | Mars_Final | Mars Final Contrast Sensitivity Score | Sensory |
| 25 | Emotion_Task_Face_Acc | Emotion Task FACE accuracy | In-Scanner Task Performance |
| 26 | Language_Task_Math_Avg_Difficulty_Level | Language Task MATH difficulty level | In-Scanner Task Performance |
| 27 | Language_Task_Story_Avg_Difficulty_Level | Language Task STORY difficulty level | In-Scanner Task Performance |
| 28 | Relational_Task_Acc | Relational Task OVERALL accuracy | In-Scanner Task Performance |
| 29 | Social_Task_Perc_Random | Social Task Overall Percentage 'Random' | In-Scanner Task Performance |
| 30 | Social_Task_Perc_TOM | Social Task Overall Percentage 'TOM' | In-Scanner Task Performance |
| 31 | WM_Task_Acc | Working Memory Task Overall Accuracy | In-Scanner Task Performance |
| 32 | NEOFAC_A | NEO-FFI Agreeableness (NEOFAC_A) | Personality |
| 33 | NEOFAC_O | NEO-FFI Openness to Experience (NEOFAC_O) | Personality |
| 34 | NEOFAC_C | NEO-FFI Conscientiousness (NEOFAC_C) | Personality |
| 35 | NEOFAC_N | NEO-FFI Neuroticism (NEOFAC_N) | Personality |
| 36 | NEOFAC_E | NEO-FFI Extraversion (NEOFAC_E) | Personality |
| 37 | ER40_CR | Penn Emotion Recognition Test: Number of Correct Responses (ER40_CR) | Emotion |
| 38 | ER40ANG | Penn Emotion Recognition Test: Number of Correct Anger Identifications (ER40ANG) | Emotion |
| 39 | ER40FEAR | Penn Emotion Recognition Test: Number of Correct Fear Identifications (ER40FEAR) | Emotion |
| 40 | ER40HAP | Penn Emotion Recognition Test: Number of Correct Happy Identifications (ER40HAP) | Emotion |
| 41 | ER40NOE | Penn Emotion Recognition Test: Number of Correct Neutral Identifications (ER40NOE) | Emotion |
| 42 | ER40SAD | Penn Emotion Recognition Test: Number of Correct Sad Identifications (ER40SAD) | Emotion |
| 43 | AngAffect_Unadj | NIH Toolbox Anger-Affect Survey: Unadjusted Scale Score | Emotion |
| 44 | AngHostil_Unadj | NIH Toolbox Anger-Hostility Survey: Unadjusted Scale Score | Emotion |
| 45 | AngAggr_Unadj | NIH Toolbox Anger-Physical Aggression Survey: Unadjusted Scale Score | Emotion |
| 46 | FearAffect_Unadj | NIH Toolbox Fear-Affect Survey: Unadjusted Scale Score | Emotion |
| 47 | FearSomat_Unadj | NIH Toolbox Fear-Somatic Arousal Survey: Unadjusted Scale Score | Emotion |
| 48 | Sadness_Unadj | NIH Toolbox Sadness Survey: Unadjusted Scale Score | Emotion |
| 49 | LifeSatisf_Unadj | NIH Toolbox General Life Satisfaction Survey: Unadjusted Scale Score | Emotion |
| 50 | MeanPurp_Unadj | NIH Toolbox Meaning and Purpose Survey: Unadjusted Scale Score | Emotion |
| 51 | PosAffect_Unadj | NIH Toolbox Positive Affect Survey: Unadjusted Scale Score | Emotion |
| 52 | Friendship_Unadj | NIH Toolbox Friendship Survey: Unadjusted Scale Score | Emotion |
| 53 | Loneliness_Unadj | NIH Toolbox Loneliness Survey: Unadjusted Scale Score | Emotion |
| 54 | PercHostil_Unadj | NIH Toolbox Perceived Hostility Survey: Unadjusted Scale Score | Emotion |
| 55 | PercReject_Unadj | NIH Toolbox Perceived Rejection Survey: Unadjusted Scale Score | Emotion |
| 56 | EmotSupp_Unadj | NIH Toolbox Emotional Support Survey: Unadjusted Scale Score | Emotion |
| 57 | InstruSupp_Unadj | NIH Toolbox Instrumental Support Survey: Unadjusted Scale Score | Emotion |
| 58 | PercStress_Unadj | NIH Toolbox Perceived Stress Survey: Unadjusted Scale Score | Emotion |
| 59 | SelfEff_Unadj | NIH Toolbox Self-Efficacy Survey: Unadjusted Scale Score | Emotion |

Supplementary Table 2: Comparison of prediction results between dynamic mode decomposition (DMD) and temporal independent component analysis (ICA). Prediction accuracy (root mean squared error, RMSE) for each of the 59 behavioral measures is presented. Asterisks indicate prediction accuracies that were significant in both the correlation and RMSE ($q \leq 0.05$).

| | DMD RMSE | DMD ($q \leq 0.05$) | ICA RMSE | ICA ($q \leq 0.05$) |
|---|---|---|---|---|
| CardSort_Unadj | 9.85 | * | 11.8 | |
| Flanker_Unadj | 9.95 | * | 11.7 | |
| Emotion_Task_Face_Acc | 3.71 | * | 4.39 | |
| CogFluidComp_Unadj | 10.7 | * | 13 | |
| WM_Task_Acc | 7.89 | * | 8.9 | * |
| ReadEng_Unadj | 9.13 | * | 10.3 | * |
| ER40ANG | 1.01 | * | 1.18 | |
| NEOFAC_E | 6.03 | | 6.8 | * |
| DDisc_AUC_40K | 0.275 | | 0.321 | |
| PicSeq_Unadj | 12.9 | | 15.2 | |
| PicVocab_Unadj | 7.9 | | 8.95 | |
| NEOFAC_O | 6.15 | | 7.36 | |
| GaitSpeed_Comp | 0.198 | | 0.23 | |
| FearSomat_Unadj | 8.1 | | 9.39 | |
| Language_Task_Story_Avg_Difficulty_Level | 1.37 | | 1.57 | |
| Strength_Unadj | 7.58 | | 8.74 | |
| Endurance_Unadj | 10.8 | | 12.5 | |
| Odor_Unadj | 9.18 | | 10.3 | |
| Dexterity_Unadj | 10.4 | | 12.1 | |
| MeanPurp_Unadj | 8.73 | | 10.6 | |
| PSQI_Score | 2.69 | | 3.13 | |
| VSPLOT_TC | 4.17 | | 4.83 | |
| ListSort_Unadj | 11 | | 12.6 | |
| PMAT24_A_CR | 4.26 | | 5.07 | |
| Relational_Task_Acc | 11.6 | | 13.6 | |
| ProcSpeed_Unadj | 15.3 | | 17.4 | |
| ER40_CR | 2.51 | | 2.8 | |
| Social_Task_Perc_TOM | 8.16 | | 9.35 | |
| FearAffect_Unadj | 7.82 | | 8.89 | |
| SCPT_SEN | 0.0792 | | 0.0885 | |
| ER40FEAR | 1.16 | | 1.38 | |
| Social_Task_Perc_Random | 10.3 | | 12 | |
| Taste_Unadj | 14 | | 16.1 | |
| Sadness_Unadj | 7.85 | | 9.36 | |
| Language_Task_Math_Avg_Difficulty_Level | 0.603 | | 0.731 | |
| Mars_Final | 0.472 | | 0.683 | |
| ER40NOE | 1.21 | | 1.41 | |
| MMSE_Score | 0.91 | | 1.03 | |
| AngHostil_Unadj | 8.54 | | 10.1 | |
| NEOFAC_C | 5.9 | | 6.9 | |
| SelfEff_Unadj | 8.46 | | 10.2 | |
| LifeSatisf_Unadj | 8.98 | | 10.6 | |
| NEOFAC_N | 7.53 | | 8.76 | |
| ER40SAD | 1.17 | | 1.37 | |
| PosAffect_Unadj | 7.96 | | 9.28 | |
| IWRD_TOT | 2.86 | | 3.21 | |
| AngAffect_Unadj | 8.23 | | 9.89 | |
| PercStress_Unadj | 9.37 | | 11.3 | |
| NEOFAC_A | 5.61 | | 6.7 | |
| ER40HAP | 0.245 | | 0.274 | |
| AngAggr_Unadj | 8.41 | | 10.1 | |
| Friendship_Unadj | 9.28 | | 11.2 | |
| PainInterf_Tscore | 7.65 | | 9.11 | |
| PercHostil_Unadj | 8.63 | | 10.2 | |
| EmotSupp_Unadj | 9.47 | | 11.4 | |
| SCPT_SPEC | 0.0343 | | 0.0396 | |
| InstruSupp_Unadj | 9.25 | | 11.2 | |
| Loneliness_Unadj | 8.75 | | 10.2 | |
| PercReject_Unadj | 8.9 | | 10.6 | |

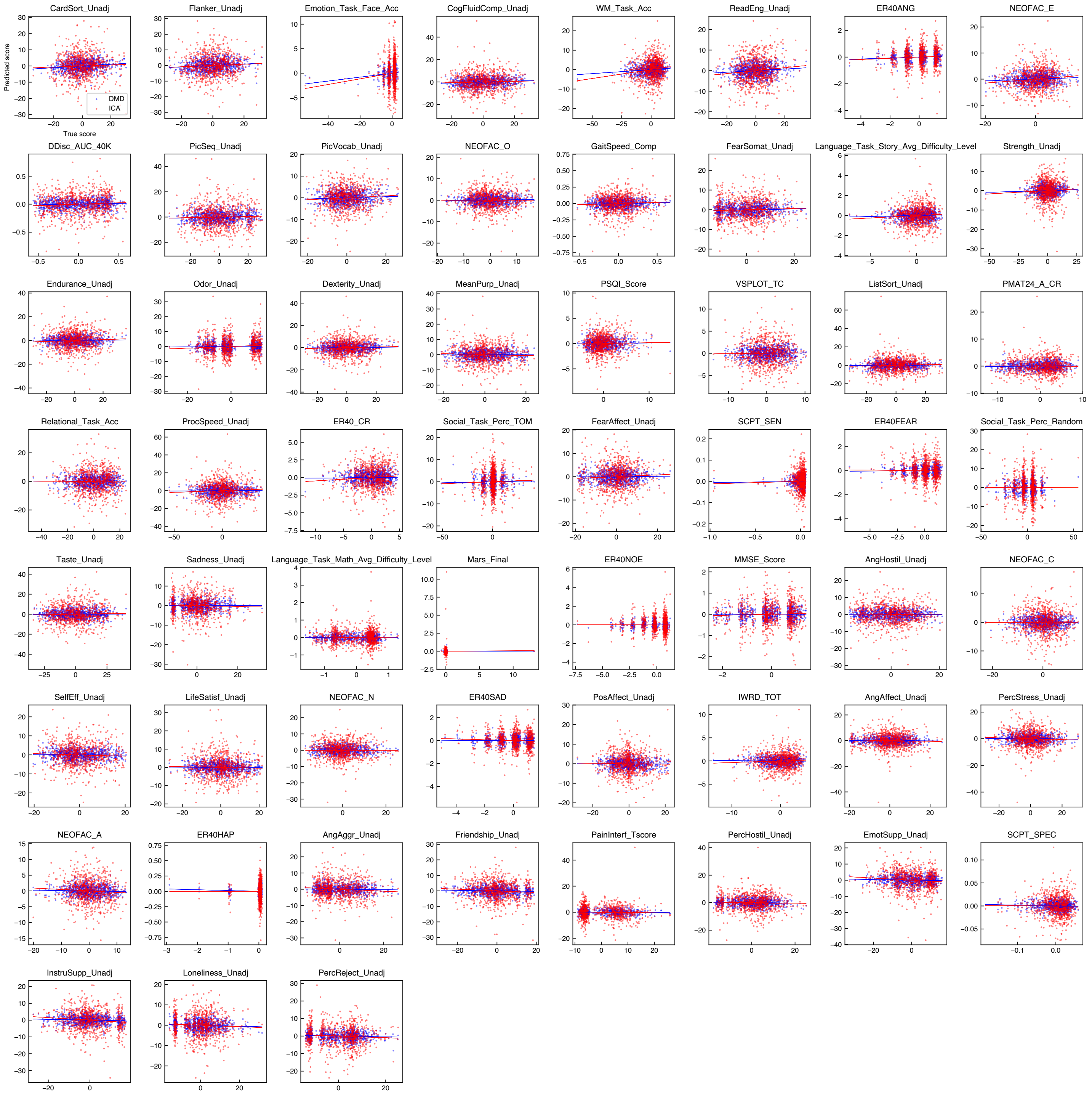


Supplementary Figure 1: Prediction results obtained using dynamic mode decomposition (DMD) and temporal independent component analysis (ICA). Individual scatter plots show associations between predicted and true behavioral scores in each of the 59 behavioral measures. Individual dots represent one subject. Each solid line shows a regression line estimated by ordinary least-squares regression.

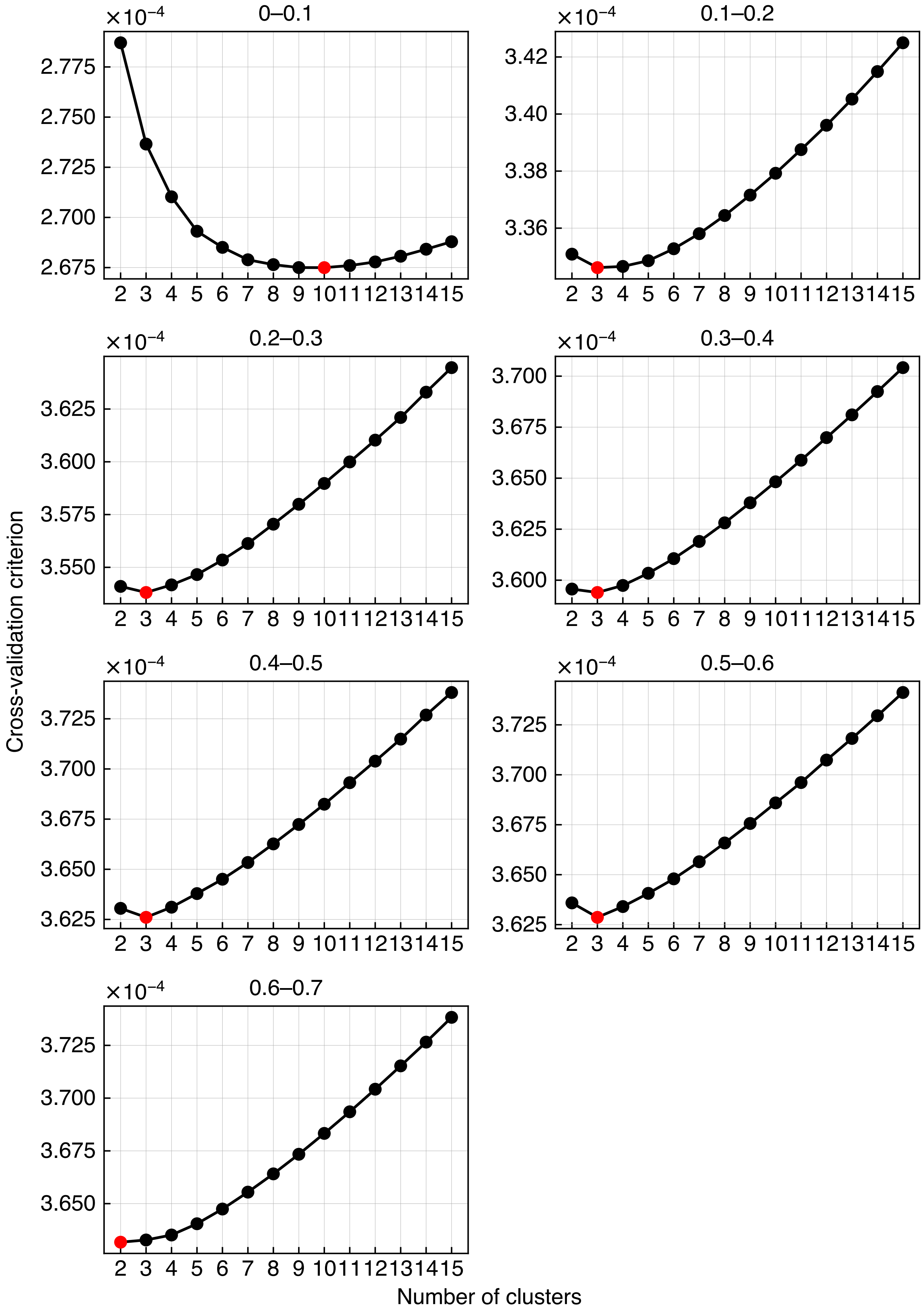


Supplementary Figure 2: Cross-validation criterion values for individual specific frequency bands. The cross-validation criterion values were computed by varying the number of clusters from 2 to 15. An optimum number was colored red.

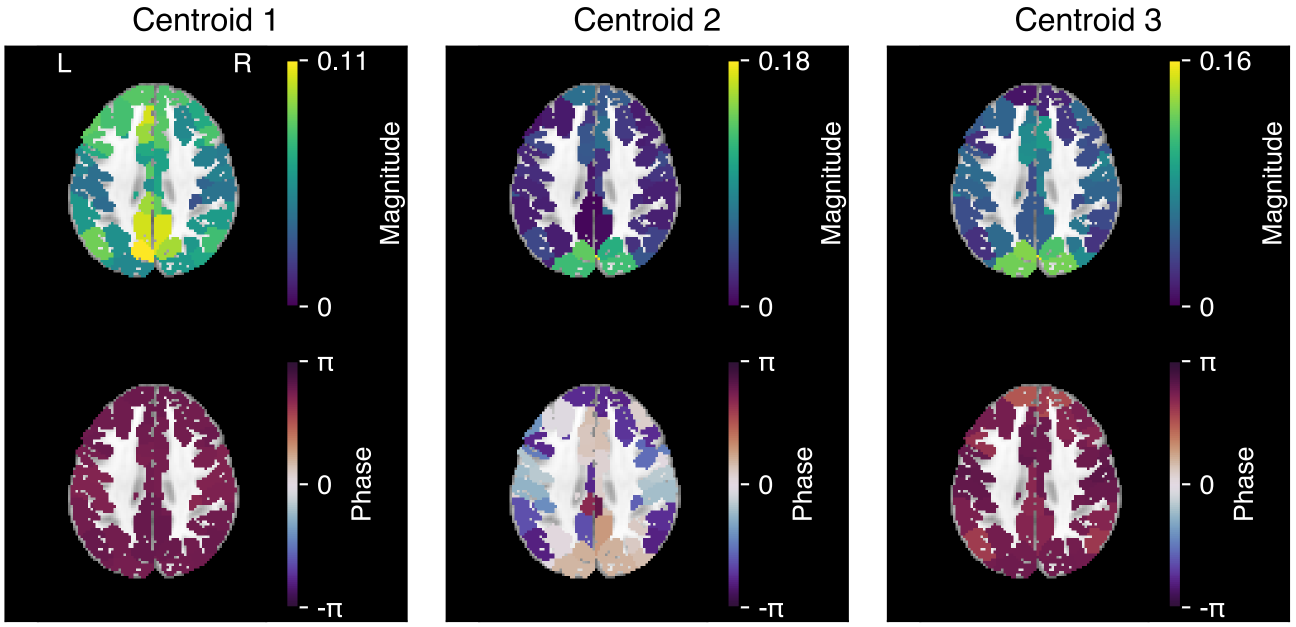


(a) Magnitude and phase maps

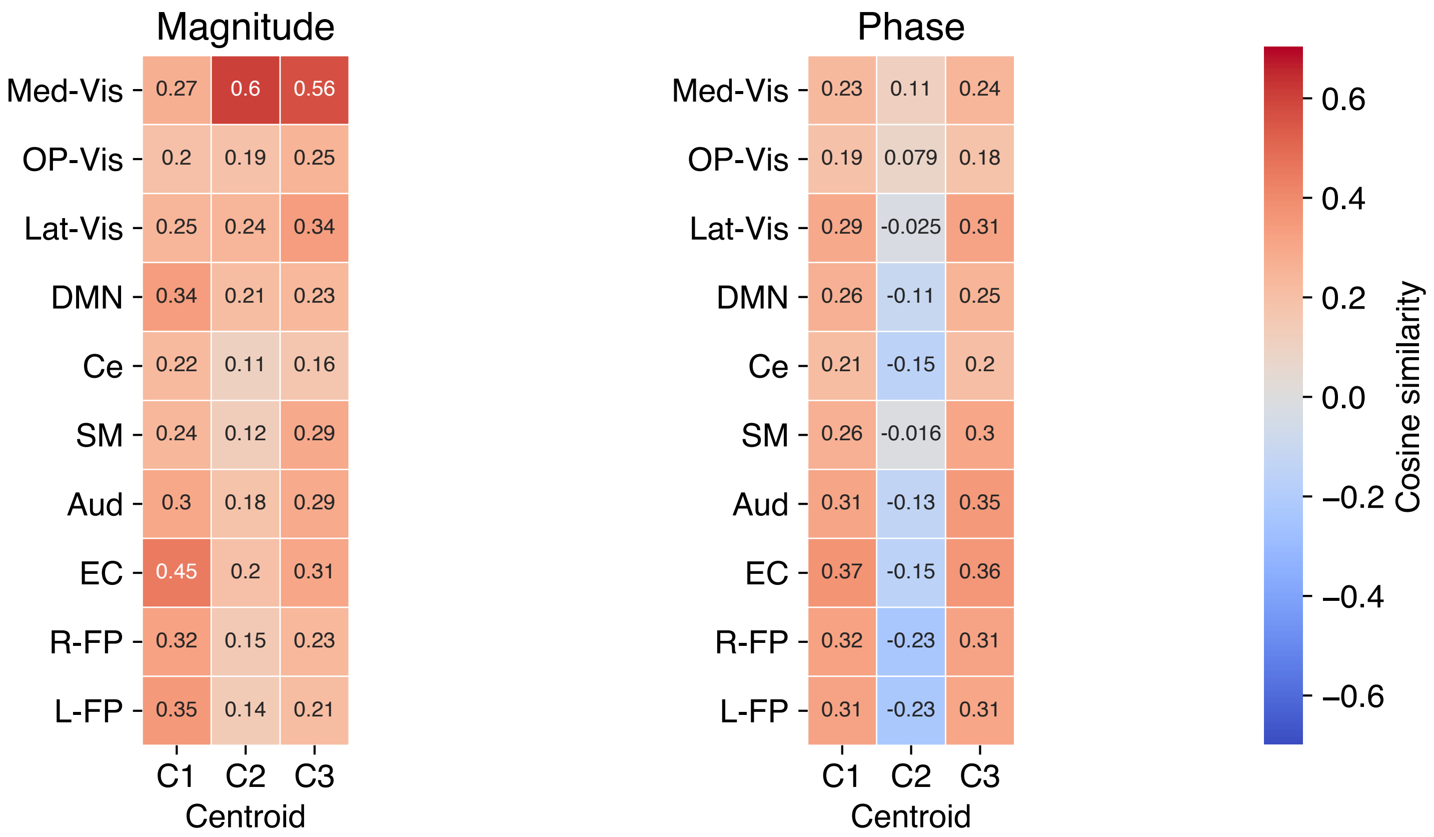


(b) Cosine similarity

Supplementary Figure 3: Clustering results (0.1–0.2 Hz). (a) Magnitude and phase maps of 3 centroids. An axial image (z = 38) for each map is shown superimposed on the skull-stripped version of the Montreal Neurological Institute (MNI152) standard space template image. (b) Cosine similarities between magnitude maps and the 10 resting-state networks (RSNs) and between phase maps and the RSNs. The 10 RSNs are as follows: the medial, occipital pole, and lateral visual areas (Med-Vis, OP-Vis, and Lat-Vis), the default mode network (DMN), the cerebellum (Ce), the sensorimotor area (SM), the auditory area (Aud), the executive control (EC), and the right and left frontoparietal areas (R-FP and L-FP).

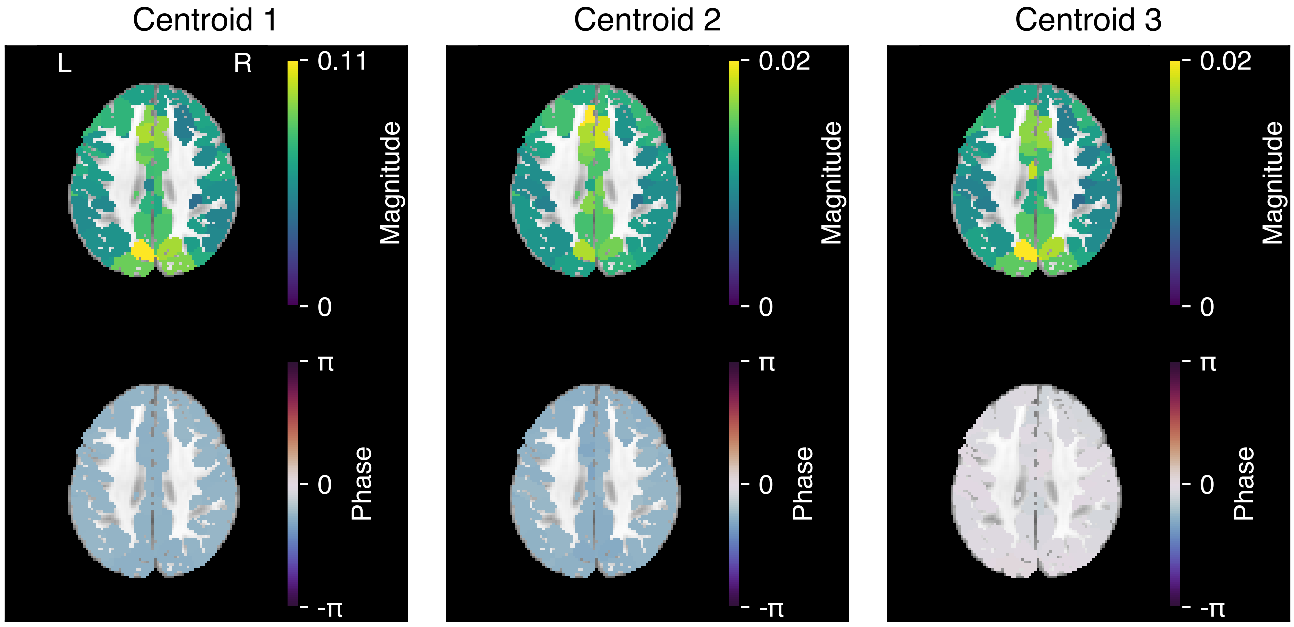


(a) Magnitude and phase maps

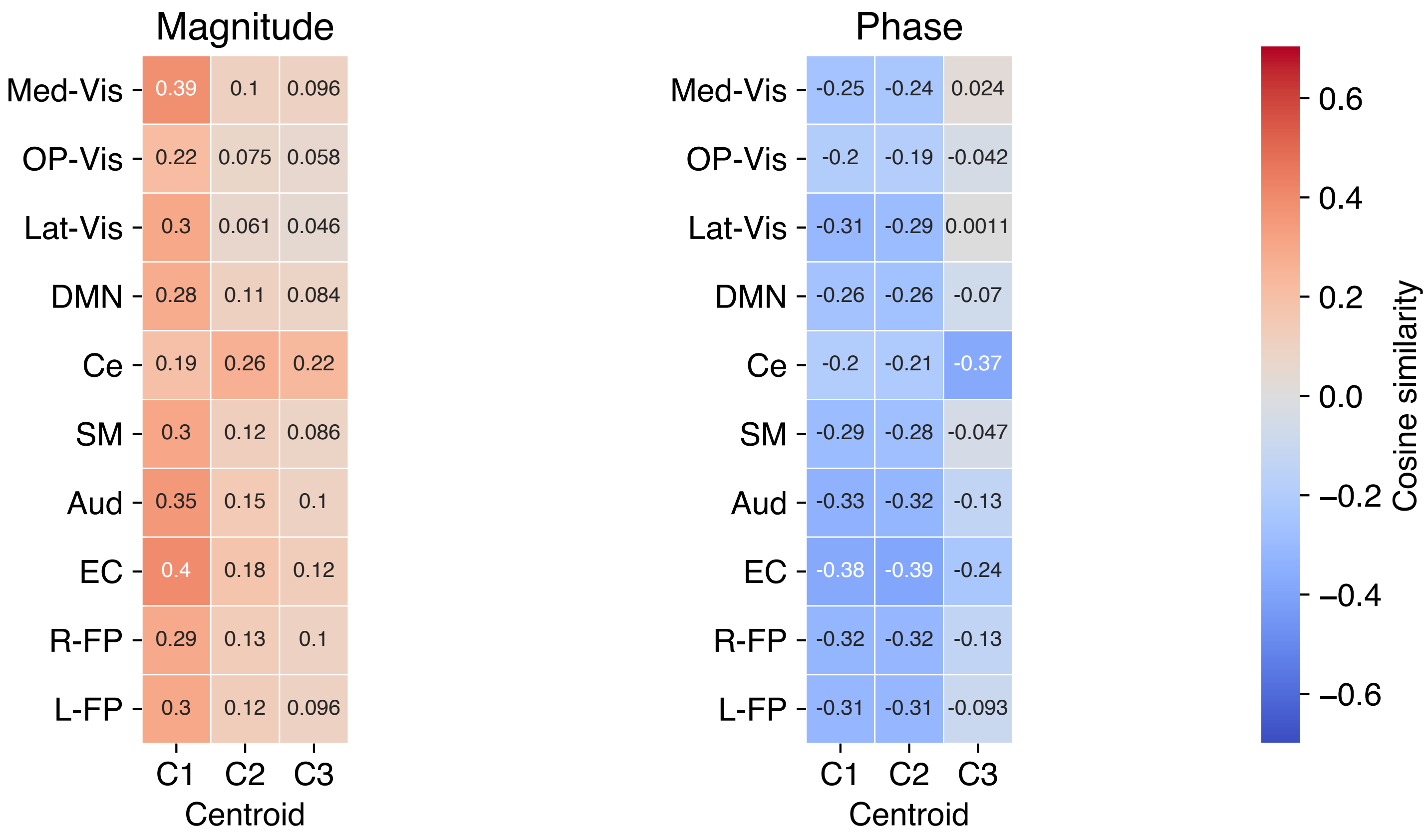


(b) Cosine similarity

Supplementary Figure 4: Clustering results (0.2–0.3 Hz). (a) Magnitude and phase maps of 3 centroids. An axial image (z = 38) for each map is shown superimposed on the skull-stripped version of the Montreal Neurological Institute (MNI152) standard space template image. (b) Cosine similarities between magnitude maps and the 10 resting-state networks (RSNs) and between phase maps and the RSNs. The 10 RSNs are as follows: the medial, occipital pole, and lateral visual areas (Med-Vis, OP-Vis, and Lat-Vis), the default mode network (DMN), the cerebellum (Ce), the sensorimotor area (SM), the auditory area (Aud), the executive control (EC), and the right and left frontoparietal areas (R-FP and L-FP).

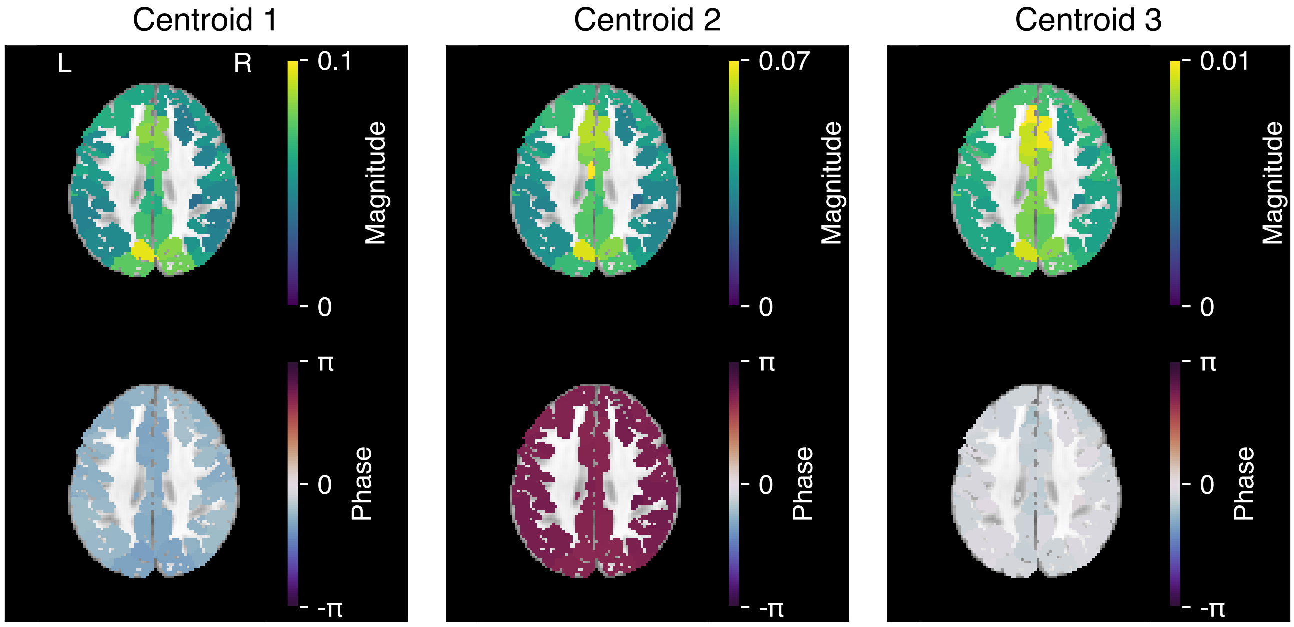


(a) Magnitude and phase maps

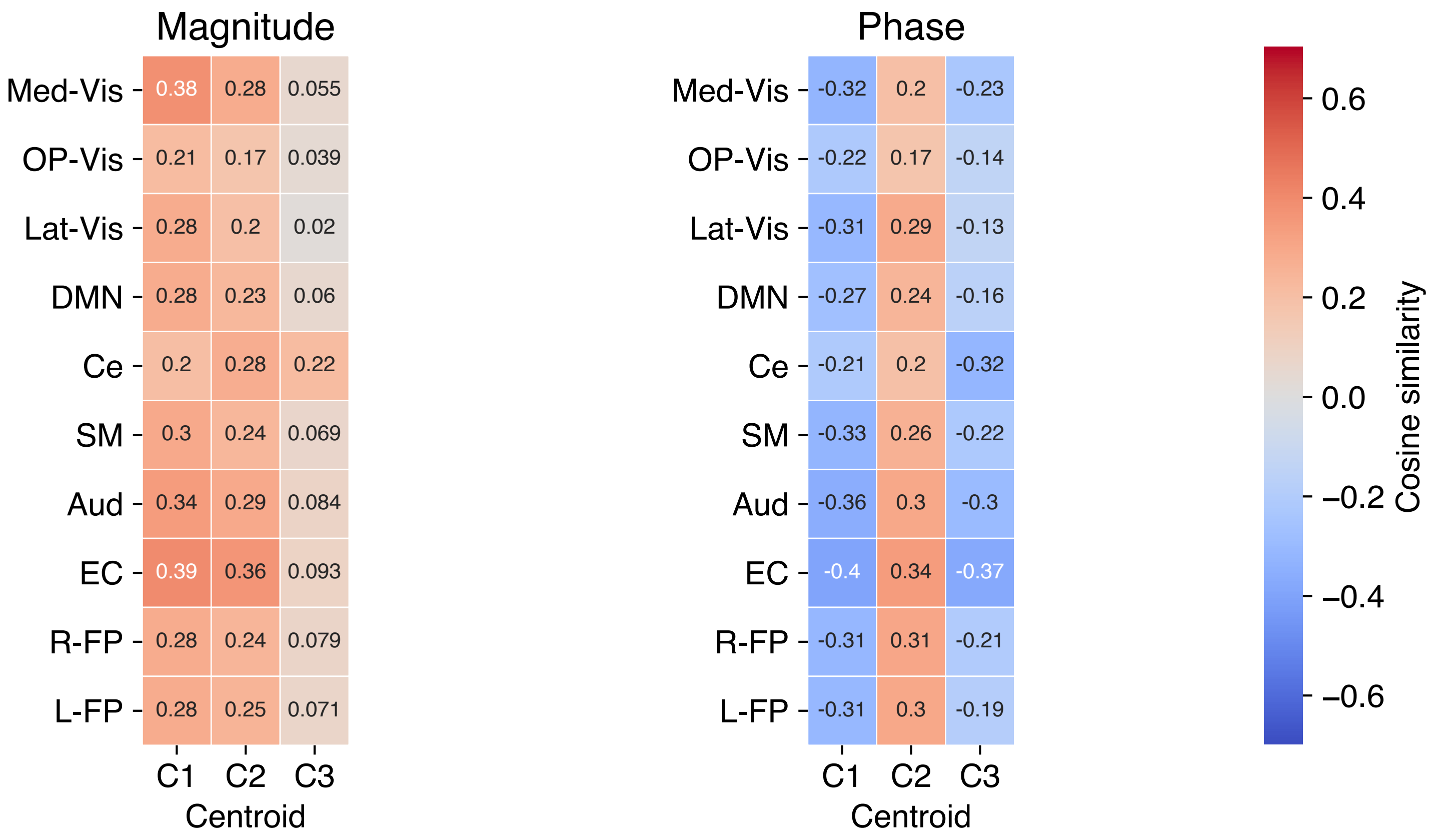


(b) Cosine similarity

Supplementary Figure 5: Clustering results (0.3–0.4 Hz). (a) Magnitude and phase maps of 3 centroids. An axial image (z = 38) for each map is shown superimposed on the skull-stripped version of the Montreal Neurological Institute (MNI152) standard space template image. (b) Cosine similarities between magnitude maps and the 10 resting-state networks (RSNs) and between phase maps and the RSNs. The 10 RSNs are as follows: the medial, occipital pole, and lateral visual areas (Med-Vis, OP-Vis, and Lat-Vis), the default mode network (DMN), the cerebellum (Ce), the sensorimotor area (SM), the auditory area (Aud), the executive control (EC), and the right and left frontoparietal areas (R-FP and L-FP).

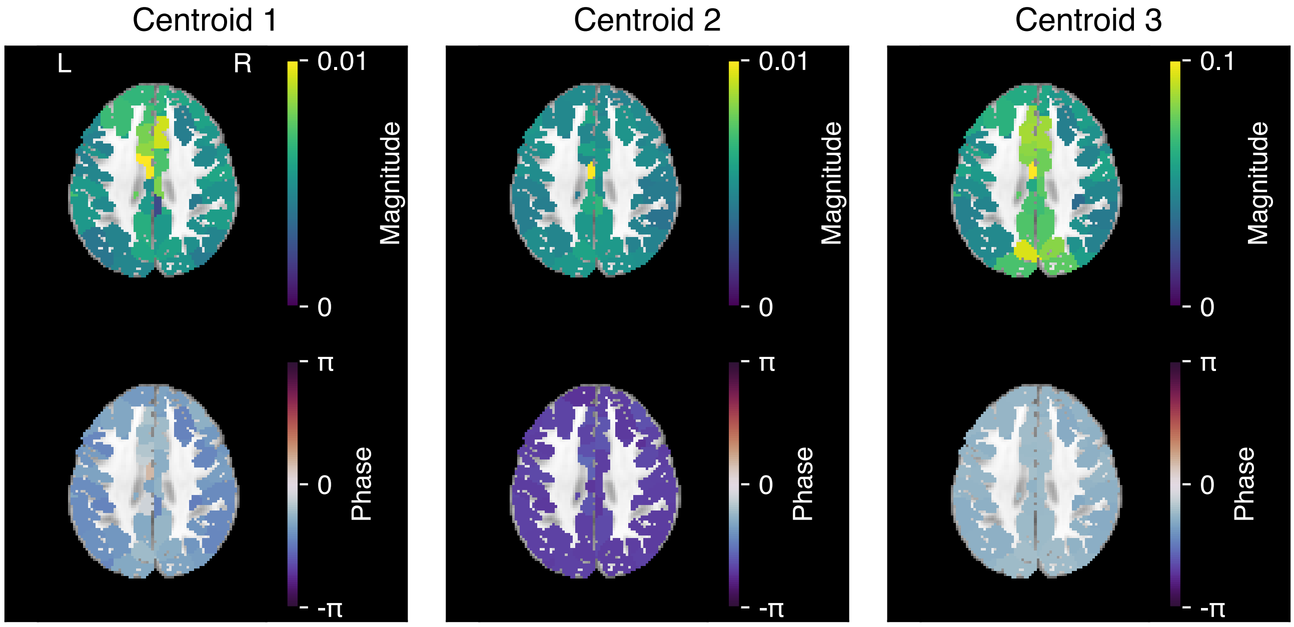


(a) Magnitude and phase maps

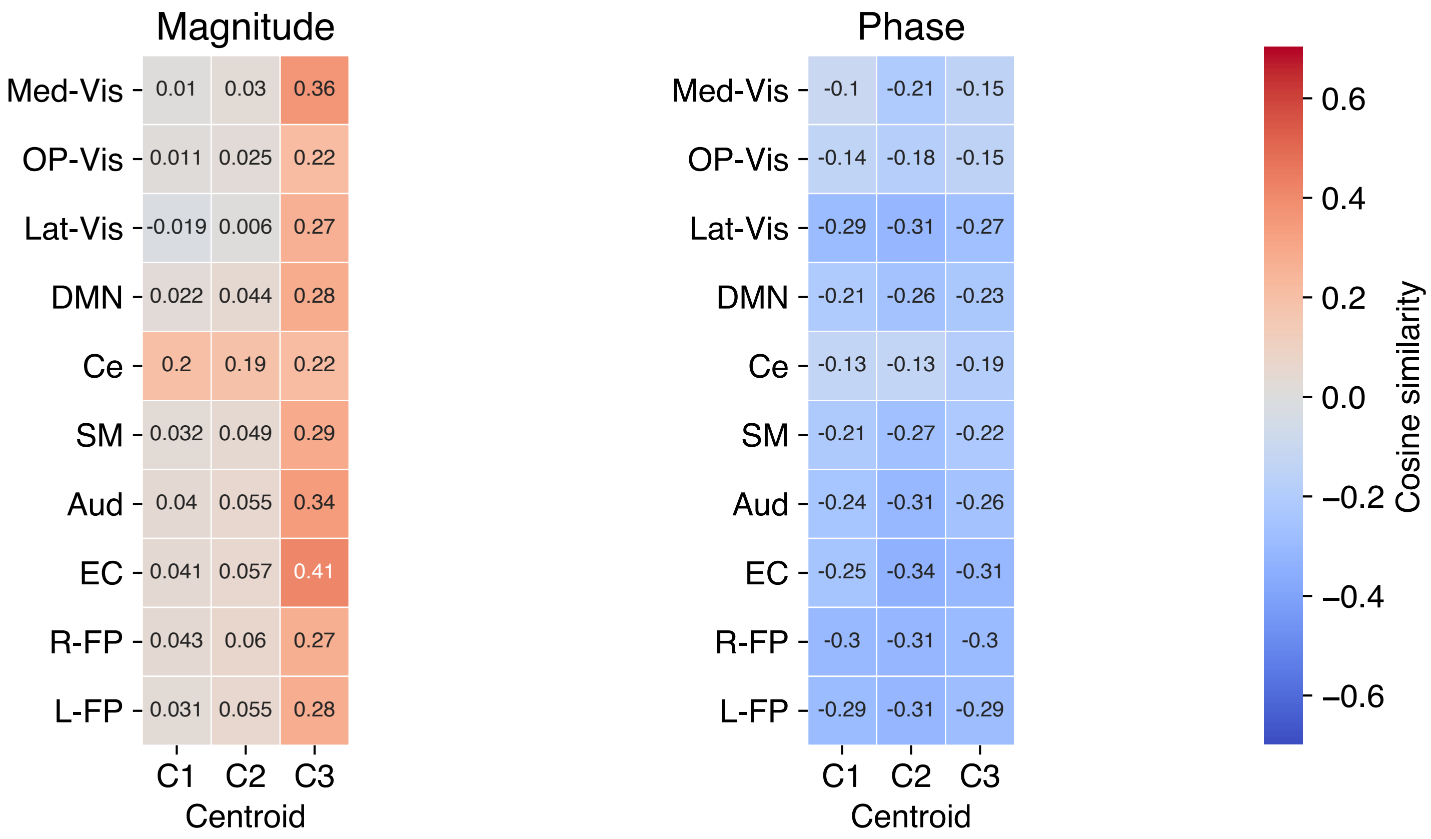


(b) Cosine similarity

Supplementary Figure 6: Clustering results (0.4–0.5 Hz). (a) Magnitude and phase maps of 3 centroids. An axial image (z = 38) for each map is shown superimposed on the skull-stripped version of the Montreal Neurological Institute (MNI152) standard space template image. (b) Cosine similarities between magnitude maps and the 10 resting-state networks (RSNs) and between phase maps and the RSNs. The 10 RSNs are as follows: the medial, occipital pole, and lateral visual areas (Med-Vis, OP-Vis, and Lat-Vis), the default mode network (DMN), the cerebellum (Ce), the sensorimotor area (SM), the auditory area (Aud), the executive control (EC), and the right and left frontoparietal areas (R-FP and L-FP).

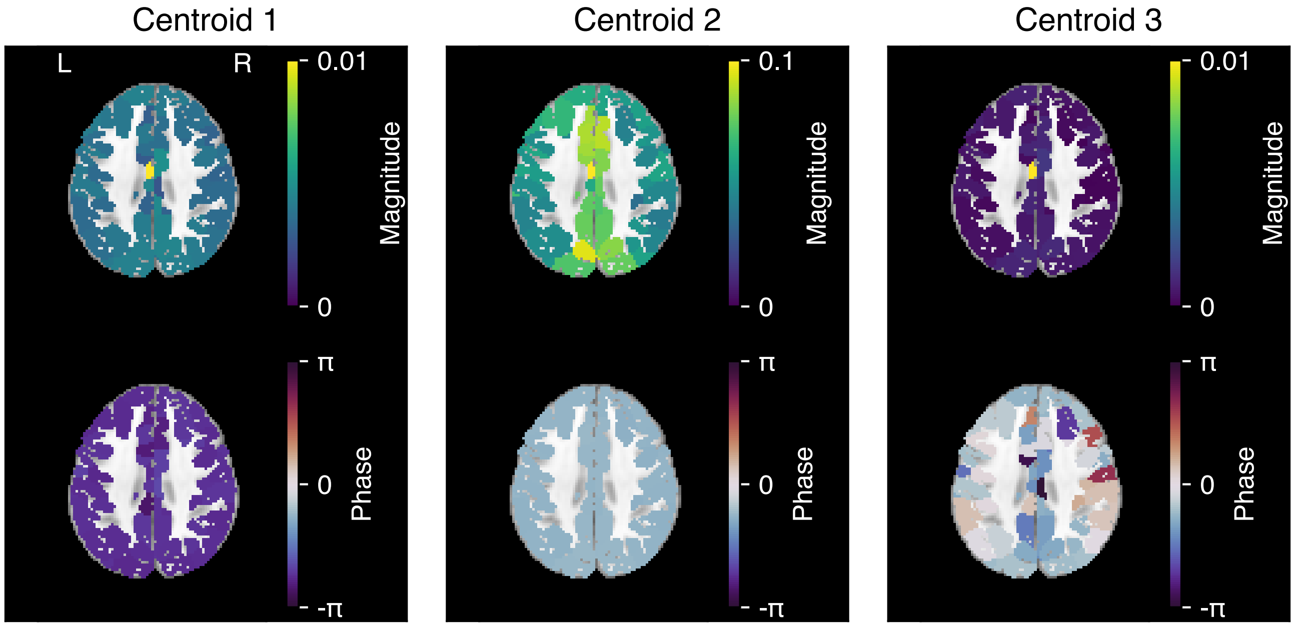


(a) Magnitude and phase maps

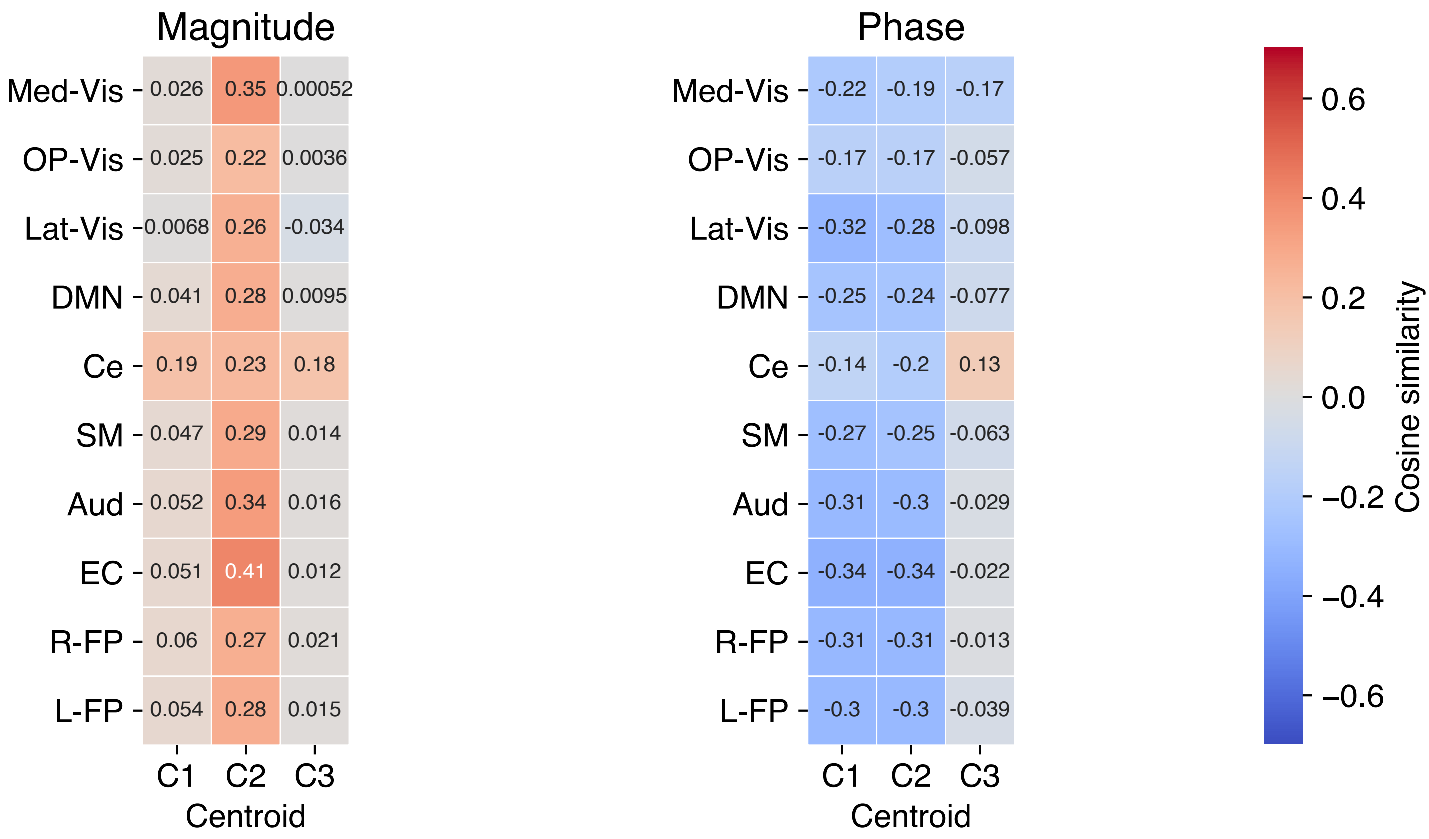


(b) Cosine similarity

Supplementary Figure 7: Clustering results (0.5–0.6 Hz). (a) Magnitude and phase maps of 3 centroids. An axial image (z = 38) for each map is shown superimposed on the skull-stripped version of the Montreal Neurological Institute (MNI152) standard space template image. (b) Cosine similarities between magnitude maps and the 10 resting-state networks (RSNs) and between phase maps and the RSNs. The 10 RSNs are as follows: the medial, occipital pole, and lateral visual areas (Med-Vis, OP-Vis, and Lat-Vis), the default mode network (DMN), the cerebellum (Ce), the sensorimotor area (SM), the auditory area (Aud), the executive control (EC), and the right and left frontoparietal areas (R-FP and L-FP).

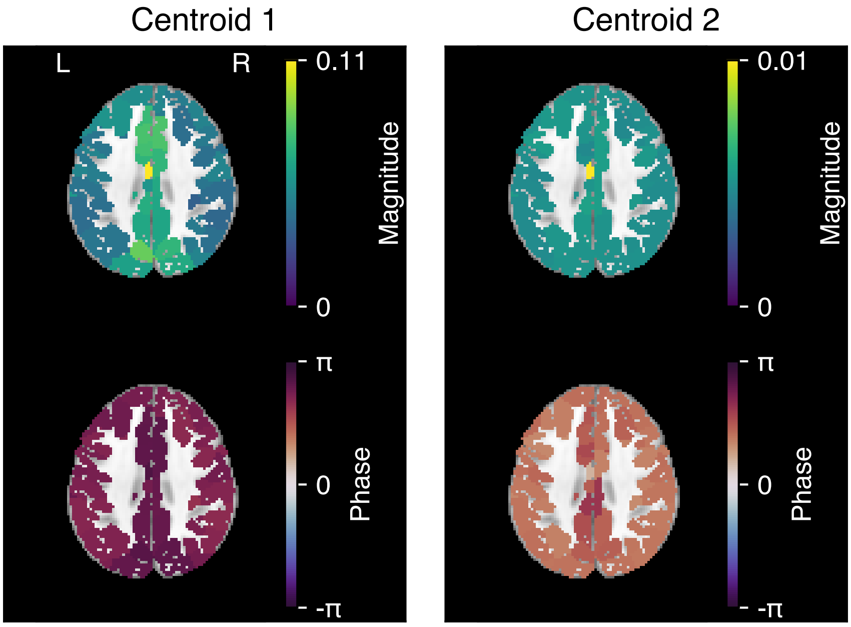


(a) Magnitude and phase maps

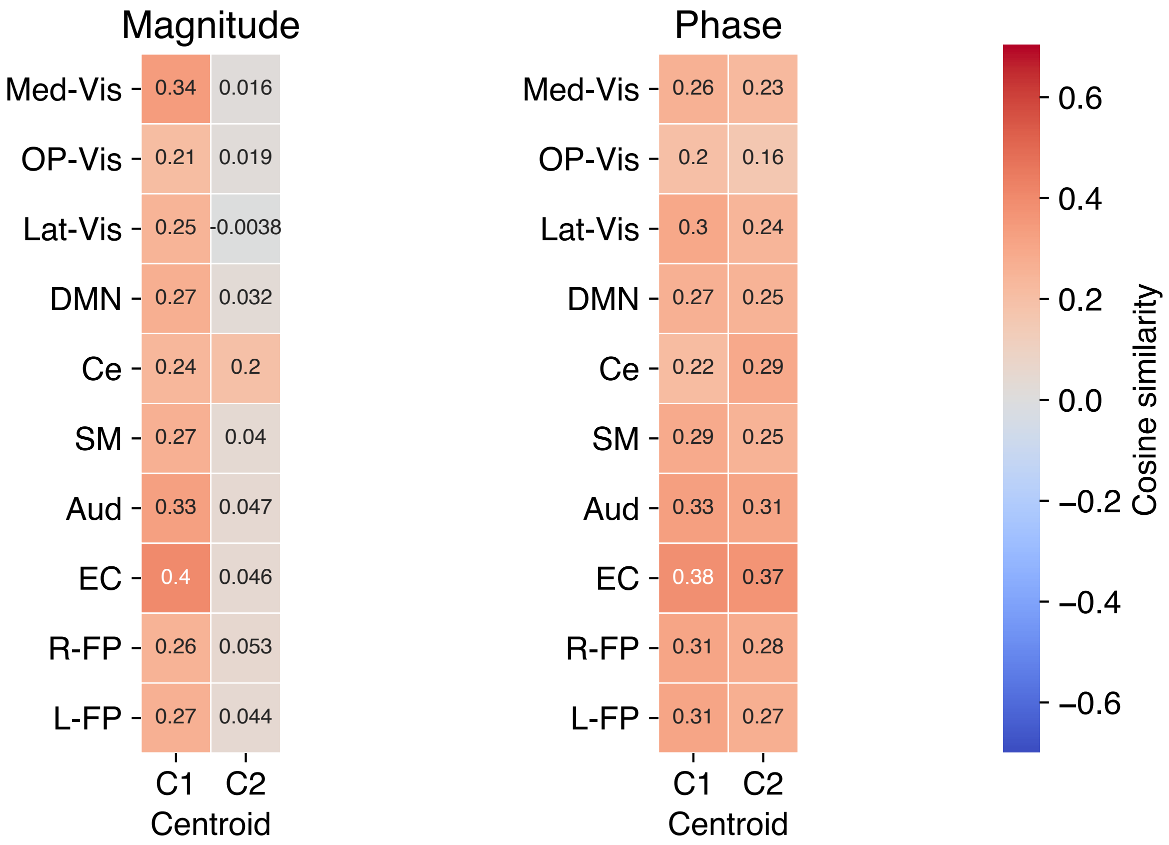


(b) Cosine similarity

Supplementary Figure 8: Clustering results (0.6–0.7 Hz). (a) Magnitude and phase maps of 2 centroids. An axial image (z = 38) for each map is shown superimposed on the skull-stripped version of the Montreal Neurological Institute (MNI152) standard space template image. (b) Cosine similarities between magnitude maps and the 10 resting-state networks (RSNs) and between phase maps and the RSNs. The 10 RSNs are as follows: the medial, occipital pole, and lateral visual areas (Med-Vis, OP-Vis, and Lat-Vis), the default mode network (DMN), the cerebellum (Ce), the sensorimotor area (SM), the auditory area (Aud), the executive control (EC), and the right and left frontoparietal areas (R-FP and L-FP).
